# Large-scale population neuroimaging reveals latent subgroup structure in functional brain organisation

**DOI:** 10.64898/2026.07.17.739129

**Authors:** S Rezvan Farahibozorg, Stephen M Smith, Lloyd T Elliott, Mark W Woolrich

## Abstract

Large-scale functional MRI datasets provide resources to understand inter-individual variation in human brain function and relate this variation to behaviour and health. However, most existing approaches fail to bridge the gap between population-average and individual-specific modelling, limiting the identification of structured subgroup heterogeneity across individuals. Here we develop a scalable framework for unsupervised subgroup discovery in population-scale resting-state fMRI data from 19,993 UK Biobank participants. Using stochastic Probabilistic Functional Modes, we estimate population-informed individualised spatial topographies of resting-state networks and derive high-dimensional functional fingerprints for each participant. We then identify latent subgroups by applying Gaussian mixture modelling independently to each fingerprint feature, yielding hundreds of reproducible subgroup definitions across 1,000 functional dimensions. We report approximately 5,700 significant differences between subgroups in a range of non-imaging phenotypes related to cognition, lifestyle, physical and mental health. Spatial organisation of the brain networks reveals distinct subgroup differences in sensory-motor and higher-order cognitive systems, in addition to correspondence with regional patterns of genetic variability across the brain. Together, these results demonstrate that large-scale functional neuroimaging contains rich latent subgroup structure linked to behavioural and biological variation. Our framework provides an interpretable and scalable basis for stratified models of human brain function and population neuroscience.

## 1 Introduction

Functional MRI provides non-invasive recordings of brain activity and connectivity. It has been instrumental in characterising various modes of human brain function, e.g., attention, language, sensorimotor, and the default mode. Each mode is a system of interconnected voxels and/or regions that work in synchrony, such that a single time course can be used to summarise their activity. Functional modes have been characterised during cognitive tasks as well as the intrinsic “resting state” brain activity (a.k.a. resting state networks, RSNs), and structured differences in brain disorders have been reported (Biswal et al., 1995; Buckner and Vincent, 2007; Calhoun et al., 2008; Damoiseaux et al., 2006; Raichle et al., 2001). With growing large-scale brain imaging datasets, e.g., Human Connectome Project (Smith et al., 2013), UK Biobank (Miller et al., 2016) and ABCD (Casey et al., 2018), neuroimaging research has been given access to open resources to examine the brain at population scales. This is fundamentally different from conventional neuroimaging datasets, where small homogeneous cohorts are available (Bijsterbosch et al., 2018; Button et al., 2013; Szucs and Ioannidis, 2020). Here, we exploit the power of big data, and the new generation of individualised brain function mapping techniques, to characterise axes of population variations and emergent subgroups based on 19,993 subjects in UK Biobank.

Brain function, and its variability, have been conventionally characterised for population averages, to understand “an average brain” (Samanez-Larkin and D’Esposito, 2008; Woolrich et al., 2025), and to compare the average brains across various cohorts, e.g., dementia patients vs age-matched healthy adults (Verdi et al., 2021). Therefore, while these studies have been important in understanding the mechanistic and consensus patterns of changes in various disorders, cross-individual variability of the brain function in health and diseases remain poorly understood. This has been partly due to the limitations of the neuroimaging datasets, where only small cohorts of < 50 people were available (Szucs and Ioannidis, 2020), and partly due to the limitation of the conventional brain connectivity modelling techniques being built on models of group averages (Calhoun et al., 2001; Yeo et al., 2011).

To address these issues, the next generation of the brain function mapping techniques have introduced individualised functional modes (Guo and Tang, 2013; Manning et al., 2018; Mejia et al., 2020). Using these new tools, growing evidence has shown that functional modes vary across individuals, and that they can be predictive of traits and diseases. However, these studies have predominantly focussed on “functional connectivity” between the modes. More specifically, after defining functional modes as either distributed systems (e.g., Default Mode or Dorsal attention), or localised parcels in atlases (e.g., Schaefer (Schaefer et al., 2018) or HCP’s Multimodal (Glasser et al., 2016) Parcellations), it has often been of interest to find temporal correlations between the timecourses of these modes, and to use these functional connectivity features as personalised fingerprints, and in phenotype prediction pipelines (Bijsterbosch et al., 2018; Finn et al., 2015; He et al., 2022).

Importantly, however, cross-individual variability in brain function is not limited to temporal features – it can also be reflected in the spatial organisation of the functional modes (Bijsterbosch et al., 2018; Harrison et al., 2020). When modelled accurately for every individual person, the spatial organisation of the modes has been shown to be associated with, and to be predictive of, cognition, emotion, lifestyle, and health. Recent studies have provided convincing evidence for this: these include hierarchical parcellation schemes that yield individualised atlases of the brain (i.e., “hard parcellations”), as well as hierarchical matrix factorisation schemes that yield individualised RSNs (i.e., “soft parcellations”) (Farahibozorg et al., 2025; Kong et al., 2019).

In this study, we build on these recent advances (**Figure 1**), in particular using stochastic Probabilistic Functional Modes (sPROFUMO) (Farahibozorg et al., 2021) on 19,993 subjects from UK Biobank to: a) characterise continuous fingerprints, i.e., axes of population variations in the spatial organisation of the brain’s functional modes; b) identify reproducible latent subgroups based on non-Gaussianities in these fingerprint features, and assess the differences between these subgroups. By combining continuous fingerprints with discrete subgroup discovery, our framework explicitly bridges population-average and individual-specific models of functional brain organisation. Importantly, we used fully unsupervised interpretable (“white-box”) techniques for both steps. First, to characterise the fingerprint features, we focussed on techniques that identify shared patterns of population variations across multiple functional modes (using linked Independent Component Analysis, FLICA) (Gong et al., 2021; Groves et al., 2011), as well as cross-subject similarities in functional organisation of the brain (subject ICA). Next, we applied Gaussian Mixture Modelling (GMM) with integrated optimisation methods to identify discrete subgroups. Rather than deriving a single clustering of the entire population, the GMM was applied to each fingerprint feature separately (i.e., 1D GMMs instead of a multivariate GMM applied to the full matrix of fingerprint features). This enables the identification of multiple subgroup definitions across different axes of functional variation, allowing distinct aspects of brain organisation to be related to different behavioural and biological characteristics.

**Figure 1.**
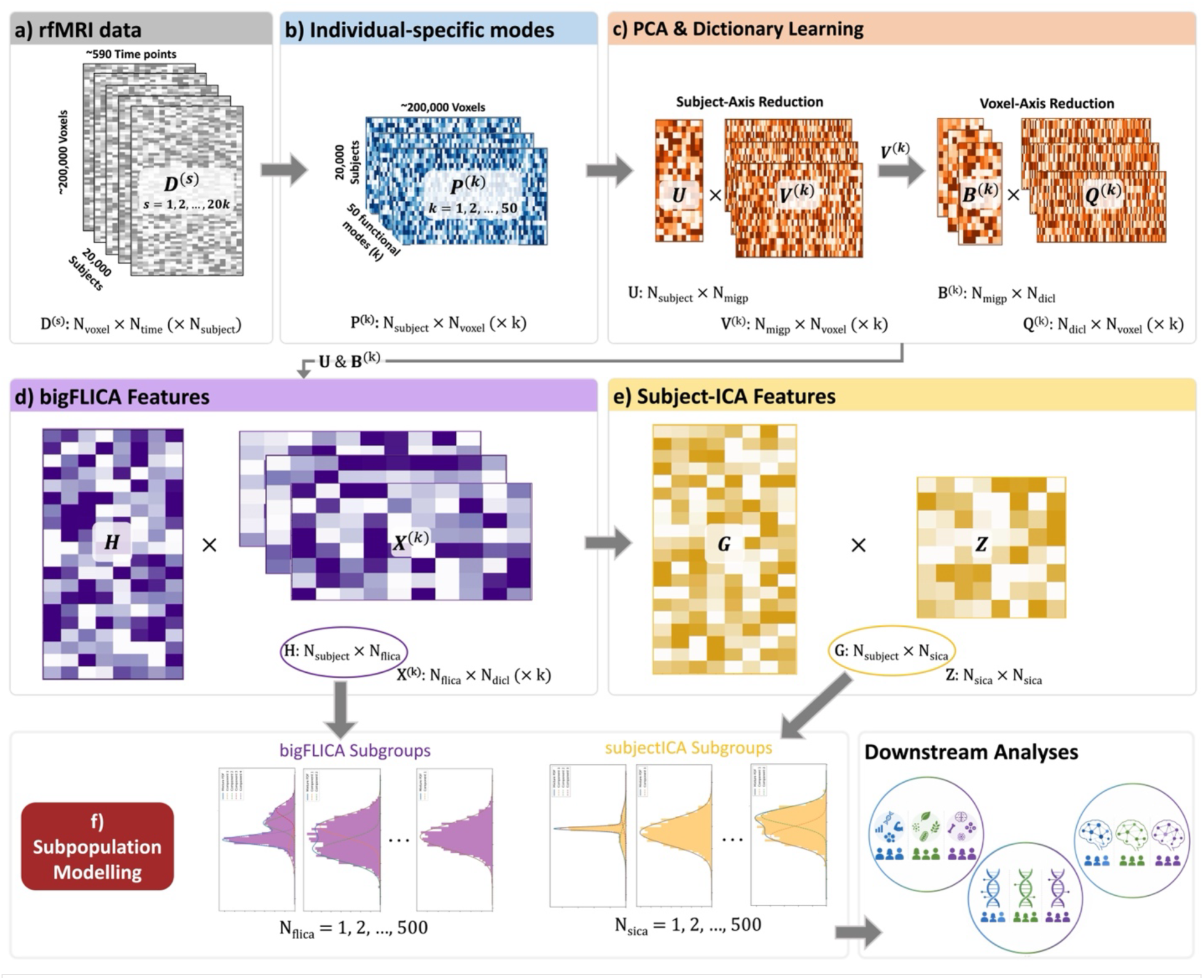
An illustration of the pipeline for subgroup identification based on functional MRI. a) Starting from preprocessed resting state fMRI (rfMRI) recordings for 19,993 UK Biobank subjects, individual-specific Probabilistic Functional Modes (PFMs) were estimated using the sPROFUMO framework. b) Spatial topographies of these modes (N_subject_ × N_voxel_ (× N_PFM_ = k)) were used as the basis for subgroup characterisation. c) PCA (multimodal MIGP with linked U estimated across PFMs) and Dictionary Learning were subsequently used for dimensionality reduction across subject and voxel axes, respectively: PCA reduced spatial maps of size N_subject_ (= 19,993) by N_voxel_ (= 228,079) to V(k) of size N_migp_ (= 1,000) by N_voxel_. Dictionary Learning further reduced V(k) matrices of size N_migp_ by N_voxel_ to B(k) of size N_migp_ by N_dicL_ (= 1,000). d) Dictionary Learning feature weightings (B(k)) were then fed into bigFLICA (a linked ICA method designed for large datasets), the output of which was weighted by PCA’s subject weightings to yield H of size N_subject_ by N_flica_ (= 500). e) bigFLICA output was in turn fed into subject ICA to characterise independent axes of subject variability, yielding G of size N_subject_ by N_sica_ (= 500). f) FLICA and subjectICA methods provided fingerprint features that were used as input to 1D Gaussian Mixture Modelling over subjects for subgroup characterisation. PFM: Probabilistic Functional Modes; PCA: Principal Component Analysis; ICA: Independent Component Analysis.

Using FLICA and subjectICA fingerprints, together consisting of 1,000 fingerprint features from which we identified 789 subgroup definitions, we show that the discovered subgroups are highly reproducible across independent splits of 19,993 UK Biobank participants. Although identified entirely without phenotype information, these subgroups exhibited widespread differences in cognition, lifestyle, physical health and mental health. Furthermore, subgroup difference maps revealed distinct spatial organisation across sensory-motor and higher-order cognitive systems. Finally, continuous fingerprint features and discrete subgroup representations showed correspondence with regional patterns of genetic variation and genes associated with neuropsychiatric disorders, health-related lifestyle and brain morphology. Together, these findings demonstrate that large-scale functional neuroimaging contains reproducible latent subgroup structure that links individual differences in functional brain organisation to behavioural and biological variation.

## 2 Materials and Methods

### 2.1 Data

#### 2.1.1 UK Biobank (UKB)

Twenty thousand (20,000) subjects were randomly drawn from the May 2019 release of the UK Biobank (UKB) data (application number 8107, all participants are over 45 years old). North West Multi-centre Research Ethics Committee (MREC) has provided ethical approval to UKB for data collection and sharing (http://www.ukbiobank.ac.uk/ethics/), and by extension, these ethical regulations cover our study. All participants completed written informed consent prior to being scanned. Resting state fMRI data in UKB consists of 1 recording session with TR = 0.735s, 490 (or 530) time points per session, providing ∼6 minutes of recording per participant. The data used here was in volumetric space and it was preprocessed using UKB’s standard preprocessing pipeline (Alfaro-Almagro et al., 2018). The preprocessing steps consisted of quality control (QC), brain extraction, motion correction, artefact rejection using FSL-FIX, and high-pass temporal filtering (Gaussian-weighted least-squares straight line fitting with sigma of 50.0s). Subsequently, data were registered to standard 2mm MNI space. According to our previous investigations into subject-specific modelling of functional modes (Farahibozorg et al., 2021), and to improve the SNR of the data, we applied an additional spatial smoothing to obtain a final 5mm FWHM smoothness (versus with the original 2mm in UKB pipeline).

### 2.2 Estimating continuous fingerprint features and discrete subgroups

#### 2.2.1 Probabilistic Functional Modes (PFMs)

Probabilistic Functional Modes (PFMs) used in this study are individualised functional modes that are obtained from applying sPROFUMO to rfMRI data from 20,000 UKB subjects. Seven subjects were removed from the subsequent analysis steps due to incomplete imaging confounds data, leaving 19,993 for the rest of the analyses. In this subsection, we present a summary of the algorithm and brief details of inference and implementation.

##### 2.2.1.1 Generative model

A detailed description can be found in our previous papers (Farahibozorg et al., 2021; Harrison et al., 2020). sPROFUMO is a hierarchical Bayesian matrix factorisation model that can be used to model functional modes in every individual simultaneously in big fMRI data. The group model provides top-down regularisation for individual specific modelling, and vice versa.

For each subject and run, fMRI timeseries (**D**^sr^) are decomposed into spatial maps (**P**s), time courses (**A**^sr^) and amplitudes (**W**^sr^), with residuals **ε**^*sr*^ :

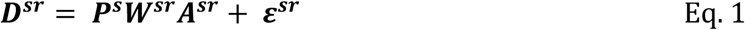

where ***D^sr^*** ∈ ℝ^***N***_*v*_×***N*_t_**^, ***P^S^*** ∈ ℝ^***N_v_***×***N_m_***^, ***W***^***sr***^ ∈ ℝ^***N_m_***×***N_m_***^, ***A***^***sr***^ ∈ ℝ^***N_m_***×***N*_*t*_**^, s stands for subject, r for scanning session. Nv, Nt and Nm denote the number of voxels, time points and modes, respectively.

The spatial maps (or just “maps”) of PFMs across the brain voxels are characterised as ***P***^*g*^. ***P***^*g*^ is estimated at the subject level with group-level map ***P***^g^ used as prior, and both ***P***^*g*^ and ***P***^*g*^ are probabilistic. A hierarchical Bayesian Double-Gaussian Mixture Model (DGMM) is used to model ***P***^*g*^ where the PFM signal and background spatial noise are each modelled using one Gaussian component of the DGMM. PFM time courses, ***A***^*sr*^, are modelled per subject and run. A multivariate normal distribution is used to model ***A***^*sr*^, and signal timecourse can be HRF-constrained, when needed (and used here). Temporal correlations between PFM timecourses (precision matrix ***α***^*sr*^) is also modelled hierarchically, and it takes the form of a Wishart distribution. HRF-constrained autocorrelations can be incorporated in the modelling of ***α***^*sr*^. PFM amplitudes, ***W***^*sr*^, which are used to capture the variance of timecourses, are modelled using a hierarchical multivariate normal distribution and are set to be diagonal. Therefore, all the PFM modelling elements are fully probabilistic, at both subject and group levels.

##### 2.2.1.2 Inference and application details

Stochastic Variational Bayes (sVB) is used as the inference technique in sPROFUMO. Specifically, during the inference, subjects from big data are divided into small batches, and model parameters are optimised to obtain an approximate distribution, *q*, that is as close as possible to the true posterior. The group model parameters, as Bayesian priors, are used for top-down regularisation of subject model parameters within each batch, and posterior distributions are summed and fed back to obtain an updated group model. Importantly, group-level parameters are treated as “global” parameters that are continuously updated over batches, and the model iterates between group and subject updates until convergence. Default parameters, as described in (Farahibozorg et al., 2021) were used for application to UKB: forget rate (*β*) = 0.6, to specify global parameter updates relying on the current compared with previous batches. Delay parameter (*τ*) = 5, to specify the contribution of the initial batches to the overall inference. Batch size of 50 subjects was used, and each subject was normally picked in 2 to 3 batches (2.5 on average), and the overall group model was updated 5,000 times.

sPROFUMO’s internal initialisation applies a few preprocessing steps to the UKB-processed rfMRI data, including voxel-wise de-meaning and variance normalisation. The first step of Bayesian model initialisation involves the estimation of a set of initial maps to start the inference from a realistic set of group-level parameters (i.e., to start the model with reasonable priors). These initial maps are estimated using MELODIC’s Incremental Group-PCA (MIGP) (Smith et al., 2014) followed by variational ICA, both of which are implemented internally within s/PROFUMO software. Subsequently, the group model is initialised using hyperpriors and the initial maps, and used for regularisation of the matrix factorisation that is conducted at subject level, as elaborated earlier, to start the full stochastic VB inference.

#### 2.2.2 FLICA and Subject ICA

Subject-specific PFM spatial maps (P) were fed into the following steps to acquire two sets of features (which together comprise the subjects’ fingerprints): FLICA features (H) and subjectICA features (G). Conceptually, FLICA finds spatially-independent components (based on spatially non-Gaussian distributions) by extracting shared information across multiple PFMs, whereas Subject ICA finds subject-independent components (i.e., based on non-Gaussian distributions in the other direction – subject-wise, not spatially). As such, they are expected to provide distinct bases for subsequent subgroup modelling.

##### 2.2.2.1 Multimodal MELODIC’s Incremental Group-PCA decomposition

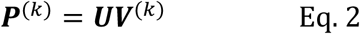

where ***P***^(*k*)^ ∈ ℝ^*N_s_*×*N_v_*^ represents the spatial maps for a mode *k*, ***U*** ∈ ℝ^*N_s_*×*N_migp_*^ represents Principal Components, and ***V***^(*k*)^ ∈ ℝ^*N*_*migp*_×*N_v_*^ represents MIGP spatial maps that are fed into the subsequent sparse dictionary learning decomposition. *N_migp_* = 1,000, *N*_*s*_ = 19,993 and *N_v_* = 228,079.

##### 2.2.2.2 Sparse Dictionary learning

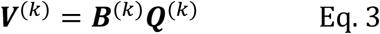

where ***B***^(***k***)^ ∈ ℝ^*N_migp_*×*N_dicL_*^ represents feature loading, and ***Q***^(***k***)^ ∈ ℝ^*N_disL_*×*N_v_*^ is the sparse spatial dictionary basis. *N_migp_* = 1,000, *N_disL_* = 1,000 and *N_v_* = 228,079.

##### 2.2.2.3 bigFLICA

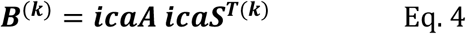

where *icaA* ∈ ℝ^*N_migp_*×*N_flica_*^ represents the mixing matrix and ***icaS*^T^** ∈ ℝ^*N_flica_*×*N_dicL_*^ represents the independent components. *N*_*flica*_ = 500.

##### 2.2.2.4 BigFLICA fingerprint features

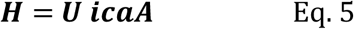

where **U** and **icaA** are derived from steps 2.2.2.1 and 2.2.2.3, respectively, yielding *H* ∈ ℝ^*N_s_*×*N_flica_*^. These are referred to as *FLICA fingerprint features* in the following sections. The FLICA fingerprint features H are subsequently used as the input to SubjectICA:

##### 2.2.2.5 SubjectICA fingerprint features

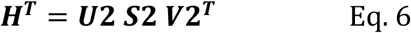

where **H** is derived from step 2.2.2.4, and further decomposed/whitened using SVD into ***U*2** ∈ ℝ^*N_flica_*×*N_sica_*^, ***S*2** ∈ ℝ^*N_sica_*×*N_sica_*^ and ***V***2^***T***^ ∈ ℝ^*N_sica_*×*N_s_*^. Next, ICA is applied to ***V*2** to obtain

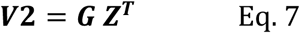

Where ***G*** ∈ ℝ^*N_s_*×*N_sica_*^, ***Z***^*T*^ ∈ ℝ^*N_sica_*×*N_sica_*^. **G** is referred to as *subjectICA fingerprint features* in the following sections. *N*_*sica*_ = 500 in this paper.

#### 2.2.3 Subgroup estimation using Gaussian Mixture Modelling

Features obtained from FLICA (H) and subjectICA (G) were fed into subgroup modelling steps independently. Here we describe the process using FLICA (H) notation; but note there is an equivalent equation using subjectICA (G) notation:

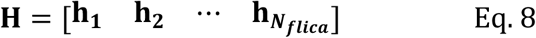

where *H* ∈ ℝ^*N_s_*×*N_flica_*^ and ***h***_*j*_ = (*h*_1*j*_, *h*_2*j*_, …, *h_*N_s_*j_*)^T^ are the N_s_ subject weights for (FLICA) feature j.

##### 2.2.3.1 Gaussian Mixture Modelling (GMM, per column)

The distribution of each column ***h****_j_* is modelled using a GMM with *K*_j_ subcomponents (i.e., subgroups):

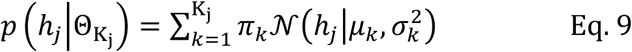

where *π*_*k*_ represents the mixing weight for the kth component (∑*π*_*k*_ = 1). *N*(·) represents Gaussian distribution with mean *µ*_*k*_ and variance 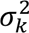. Θ_*K*_ represents the set of model parameters {(*π*_*k*_, *µ*_*k*_, 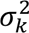) ∣ *k* = 1, …, *K*_j_}.

##### 2.2.3.2 Optimisation and Model Selection

The optimal number of subcomponents, 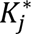, is selected from *K*_j_ ∈ {1, 2, …, 20} based on two criteria, Bayesian Information Criterion (BIC) and Davies-Bouldin Index (DBI), as follows:

###### a) Bayesian Information Criterion (BIC)

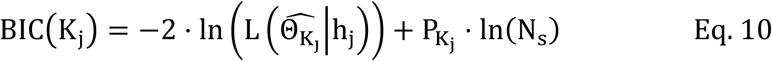

where 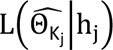 is the maximised likelihood of the *K_j_*-component model given the data. *P_K_j__* is the number of free parameters (3K_j_ - 1), and *N*_*s*_ is the number of subjects. The optimal number 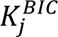 was selected as the number of subcomponents that minimised the BIC:

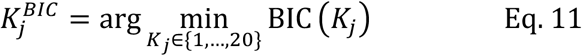

###### b) Davies-Bouldin Index (DBI)

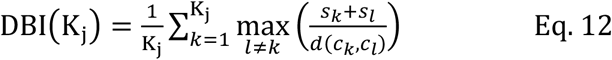

where *s*_*k*_ is the average distance within cluster *k* (intra-cluster scatter), and *d*(*c*_*k*_, *c*_*l*_) is the distance between centroids *c*_*k*_ and *c*_*l*_ (inter-cluster separation). The optimal number 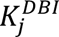 was selected as the number of subcomponents that minimised the DBI (DBI = 0 is the optimal score):

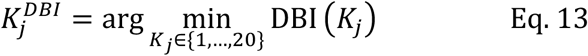

The final number of subgroups 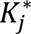 for component *j* was selected as the minimum of the two optimal numbers: 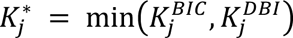. This was repeated for j = 1, 2, …, *N*_*flica*_. The same process was applied to columns of G, the subject ICA features, j = 1, 2, …, *N*_*sica*_.

This approach yields the optimal number of subgroups from a mathematical point of view. However, for example, with super-Gaussian distributions that have long tails at both ends, the GMM would typically merge both tails as one Gaussian distribution (i.e., one subgroup). However, biolosical interpretation of the two tails is expected to be distinct: e.g., if one represents the highest end of a fingerprint feature related to fluid intelligence, the other will represent the lowest end, and thus, it is best to split the two. Therefore, in scenarios like this we split that component into two halves, each consisting of one end of the distribution.

It is worth noting that we applied a separate 1D GMM to each fingerprint feature separately (i.e., columns of H and G). This was preferred over a multivariate GMM applied to the full matrix of fingerprint features at once. The latter would provide a single definition of subgroup divisions across the entire feature space, whereas the 1D approach used here can potentially provide hundreds of distinct subgroup divisions, each possibly linked to its own specific set of phenotypes or disease. In the Results section, we use Kurtosis and Skewness to characterise non-Gaussianities in each fingerprint feature. Kurtosis = 0 indicates a Gaussian distribution while Kurtosis > 0 and < 0 would indicate super-Gaussian and sub-Gaussian distributions, respectively. Skewness = 0 indicates a Gaussian distribution while Skewness > 0 and < 0 represent long tails on the right-hand and left-hand side, respectively.

#### 2.2.4 Confound Removal in UK Biobank

Imaging confounds refer to factors such as head motion or head size, that tend to correlate with both non-imaging phenotypes (e.g., disease scores) and brain-imaging data. This can become particularly problematic in studies with big data populations (high sample sizes providing high statistical power), when associations between imaging and non-imaging variables are of interest: confounds can significantly affect the interpretability of the findings (Alfaro-Almagro et al., 2020). In this study, imaging confounds can additionally impact and bias subgroup identifications. Therefore, we deconfounded the raw subject-specific PFM spatial maps. For this purpose, **P^s^** in Eq. 1 was concatenated across subjects and deconfounded, before conducting the subsequent feature estimation (i.e., FLICA and Subject ICA, see section 2.2.2) and subgroup identifications. This was done by linearly regressing out the confounds from the PFM spatial maps, and where relevant, the non-imaging variables. The list of imaging confounds is included in Table S 1.

#### 2.2.5 Non-imaging derived phenotypes (nIDPs)

We used 436 non-Imaging-Derived Phenotypes (nIDPs) from UK Biobank to test phenotypical differences between subgroups, and Canonical Correlation Analyses between imaging and non-imaging variables, as explained below. These nIDPs spanned six categories: cognitive, mental health, alcohol, tobacco, cardiovascular and bone health.

##### Cognitive nIDPs

68 out of 1172 nIDPs related to cognitive tests were selected based on two criteria: firstly, we manually prefiltered the outcome measures, to only include active metrics of subjects’ performance. For example, under the “Reaction Time” cognitive test, which consists of viewing two cards (A and B) and pressing buttons when identical, “Mean time to correctly identify matches”, was labelled as active (included) whereas “Index for card A in round” was labelled as passive (excluded). Secondly, only tests that had a non-missing value in at least 25% of the subjects were included. The final list consisted of 68 cognitive nIDPs in the following categories: Reaction time, Trail making, Matrix pattern completion, Numeric memory, Prospective memory, Pairs matching, Symbol digit substitution and Fluid intelligence. Additional details of these nIDPs can be found in Table S 2 and here: https://biobank.ctsu.ox.ac.uk/crystal/label.cgi?id=100026.

##### Mental Health nIDPs

We selected 142 metrics related to mental health that had a non-missing value in at least 25% of the subjects. Details of these nIDPs are available in Table S 3 and the following link: https://biobank.ctsu.ox.ac.uk/crystal/label.cgi?id=100060.

##### Alcohol and Tobacco nIDPs

We selected 37 Alcohol-related and 19 Tobacco-related metrics that had a non-missing value in at least 25% of the subjects. More details about these nIDPs are available in Table S 4, Table S 5 and the following links: https://biobank.ctsu.ox.ac.uk/crystal/label.cgi?id=100051 and https://biobank.ctsu.ox.ac.uk/crystal/label.cgi?id=100058.

##### Cardiovascular nIDPs

77 out of 992 nIDPs related to cardiovascular health were selected based on two criteria: firstly, we manually prefiltered the nIDPs and excluded the metrics that were not directly health-related. For example, “Systolic blood pressure, manual reading (0.0)” and “LV stroke volume (2.0)” were included but “Program category (0.0)” and “Completion status of test (0.0)” were excluded. Secondly, only tests that had a non-missing value in at least 25% of the subjects were included. Details of these nIDPs can be found in Table S 6 and the following links: https://biobank.ctsu.ox.ac.uk/crystal/label.cgi?id=100011, https://biobank.ctsu.ox.ac.uk/crystal/label.cgi?id=104 and https://biobank.ctsu.ox.ac.uk/crystal/label.cgi?id=100012.

##### Bone Health nIDPs

We selected 93 metrics related to bone health that had a non-missing value in at least 25% of the subjects. More details about these nIDPs are available in Table S 7 and the following link: https://biobank.ctsu.ox.ac.uk/crystal/label.cgi?id=100018.

#### 2.2.6 Statistical comparison of phenotypes between subgroups

As elaborated earlier in 2.2.2 and 2.2.3, FLICA and subjectICA features and the subsequent subgroup estimations were unsupervised. We next tested if the subgroups were significantly different with respect to each of the 436 phenotypes described in 2.2.5. For this purpose, for each FLICA/subjectICA feature, and each nIDP, we compared the 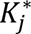 identified subgroups using the Kruskal-Wallis test, a non-parametric test that allows the comparison of two or multiple subgroups. The obtained p-values were then adjusted for multiple comparisons using both Bonferroni and (separately) FDR criteria (correcting across features x nIDPs), and results were summarised using Manhattan plots. The FLICA/subjectICA features for each nIDP that passed the Bonferroni correction threshold will be referred to as **nIDP-hits** in the rest of the paper.

#### 2.2.7 Estimating spatial Maps of Subgroups Differences (MSDs)

We next estimated the spatial Maps of Subgroup Differences (MSD) across the brain voxels. This was done independently for each resting state network (RSN). The process was as follows:

1. For each nIDP, we extracted the nIDP-hits, after p-value calculation and Bonferroni correction for multiple comparisons, as described in 2.2.6. This was done for both subjectICA and FLICA features. To focus on the strongest nIDP-hits and additionally to reduce computational costs, we extracted up to up to 10 significant fingerprint features per nIDP. For example, for “Fluid intelligence score (0.0)”, we will have N_hit_FLICA_ <= 10 and N_hit_subjectICA_ <= 10 hits.
2. 11 prominent RSNs (6 cognitive and 5 sensory-motor) were manually selected: cognitive RSNs consisted of Default Mode 1&2, left and right Fronto-parietal, Ventral Attention and Language; sensory-motor RSNs consisted of Visual 1&2, Motor 1&2 and Auditory. We analysed subject-specific PFM spatial maps (P in Eq. 1), one at a time, and concatenated the spatial maps across subjects, obtaining N_s_ x N_v_ matrices per RSN.
3. For each of the nIDP-hits, and per brain voxel, we conducted Kruskal-Wallis test across the 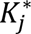 subgroups associated with that nIDP-hit, and computed p-values. These show how each RSN spatial map’s intensity at a given voxel differs across the 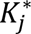 subgroups.
4. We finally conducted Bonferroni correction for multiple comparisons across (N_hit_FLICA_ + N_hit_subjectICA_) x Nphenotype_per_category. These maps were then thresholded at corrected p-value < 0.05 to obtain binarised masks of significant differences. This yielded (N_hit_FLICA_ + N_hit_subjectICA_) x N_phenotype_per_category_ binarised spatial maps.
5. Maps were summed across all N_hit_FLICA_ + N_hit_subjectICA_ phenotypes per category, normalised per category, then summed across categories, yielding MSDs.

#### 2.2.8 Canonical Correlation Analysis: multivariate associations between subgroups and traits

Canonical Correlation Analysis (CCA) was used to find multivariate mappings between each of the nIDP categories and subjectICA/FLICA fingerprints, subgroups, or both concatenated. Each CCA component reflects a weighted combination of PFM-based features and a weighted combination of nIDPs, to obtain transformed outputs that are maximally correlated. In order words, CCA projects data onto axes of population co-variation between brain and non-imaging traits. CCA used in this paper was aimed at examining the phenotypical relevance of subgroup estimations, and to test whether subgroups-alone, or the combination of subgroups+fingerprints would show stronger association to traits, than the conventional continuous feature spaces alone.

##### Conducting CCA

CCA consisted of the following steps:

1. We conducted cross-validated CCA using 5-fold cross-validation. Therefore, the Ns subjects were randomised and divided into 5 folds, and in each iteration, 4 folds were used to train the CCA, as well as any additional normalisation and dimensionality reduction steps, as described below. The remaining fold was then used as test set and results were averaged across folds for reporting.
2. One side of CCA received brain features, in either of the following three forms:

a. Fingerprint-features-only: Continuous fingerprint features obtained from FLICA and subjectICA. Columns of these matrices were concatenated, standardised, and additionally, following guidelines from (Helmer et al., 2024) regarding the impact of sample to feature ratio on CCA stability, we dimension-reduced the input feature matrices to 100 features using PCA before feeding into CCA.
b. Subgroup-only: subgroup summaries were one-hot transformed and dimension-reduced to 100 features using Multiple Correspondence Analysis (MCA), which is the PCA-equivalent for categorical data.
c. Fingerprint + Subgroup: As will be shown later in Results section 3.1, some of the FLICA/subjectICA components (i.e., fingerprint features) did not yield distinct subgroups (i.e., were inherently continuous Gaussian features). Step 2a was repeated on those features and 100 principal components were estimated. This was then concatenated with features obtained from step 2b.
3. The other side of CCA received phenotype matrices as input: Ns × NnIDP. This was done separately for the six nIDP categories: cognition, mental health, tobacco, alcohol, cardiovascular and bone health. Given the high number of missing values in UK Biobank nIDPs, we used sklearn’s SimpleImputer with mean imputation to account for the missing values. To account for any redundancies, or highly correlated columns, especially after imputation, we applied PCA to the resulting nIDP matrices before feeding into CCA (as before, all the steps are conducted within the cross-validation loop).
4. CCA was conducted for 3 (image feature spaces) by 6 (nIDP category) pair-wise comparisons, yielding 18 results overall.
5. CCA yields a linear transformation of PFM feature matrices (X) and phenotype feature matrix (Y) so as to maximise their correlation; i.e. Y*A=U ∼ X*B=V, where U and V are the linearly-transformed versions of the nIDP and PFM feature matrices, respectively.
6. For each cross-validation fold, we computed the correlation between corresponding test-set canonical variates (the diagonal of the U–V cross-correlation matrix), yielding one correlation per CCA component. Because component ordering can vary across folds, we used the first fold as a reference and matched components in subsequent folds using the Hungarian algorithm (scipy.optimize.linear_sum_assignment). Correlations were then reordered according to this matching before pooling across folds. Averaging the matched correlations provided an estimate of the shared variance captured by each CCA component pair.

#### 2.2.9 Genetic relevance of subgroups

We next examined whether spatial maps of subgroup variability in any given RSN can be linked to the spatial maps of genetic variability in that RSN. The analysis pipeline was as follows:

The first step included a Genome-Wide Association Study (GWAS) of all the FLICA/SubjectICA fingerprint features (500 + 500) to identify single nucleotide polymorphisms (SNP), i.e., **SNP-hits**, across the Genome. We used BGENIE software for GWAS analysis (Bycroft et al., 2018), and a similar processing pipeline as described in (Smith et al., 2021) (the latest 2025 software update). Before feeding into GWAS analysis, each fingerprint feature (num_subjects x 1) vector was Gaussianised using a rank-based inverse normal transformation as implemented in the FSLNets MATLAB package, deconfounded for genetic principal components, and standardised by demeaning and variance normalisation. Additional filtering of the genetic data included MAF (minor allele frequency) ≥ 0.01, imputation information scores ≥ 0.3 (for the Manhattan plots), uncorrected GWAS p-value threshold of 10−7.5, and a Bonferroni-corrected threshold of 10−10.2. Of the 19,993 subjects included in this study, ∼16,000 were included in UKB’s BIG40 (updated with 63k subjects in 2025) sample, after applying genetic and ancestry filters. These subjects were divided into Discovery (n = 11,198) and Replication (n = 5,583) cohorts, and SNP-hits were identified for both cohorts, as well as the overlap between them. After extracting the SNP-hits, we further used BGENIE to extract Allele dosage for the hits, yielding a matrix of Ns x NSNP_hit. After identifying the SNP-hits, we additionally used FUMA (Watanabe et al., 2017) to link the SNPs to genes, and the genes to functions and disease.

The second step involved estimating spatial Maps of Allele Dosage (MAD) per RSN, aiming to compare those spatial maps to the corresponding MSDs described in section 2.2.7. For this purpose, we analysed subject-specific PFM spatial maps (P in Eq. 1), one at a time, and concatenated the spatial maps across subjects, obtaining Ns x Nv matrices per RSN. Each of these were then correlated with Allele dosage maps of size Ns x NSNP_hit across subject dimension, thus yielding NSNP_hit voxel-wise MADs. Therefore, for each PFM/RSN, we estimated 1 MSD (see section 2.2.7) and NSNP_hit MADs. We computed spatial correlation between MSDs and MADs, thus obtaining a correlation matrix of size NSNP_hit x Npfm. Similar to section 2.2.7, 11 RSNs were used for these analyses.

It is worth noting that we restricted the SNP-hits used in the spatial map analyses to either: a) hits of the Discovery cohort that were supported by multiple nearby SNPs in linkage disequilibrium (LD) that exceeded the genome-wide significance threshold; these clustered peaks are more likely to represent true loci rather than technical artefacts; or b) isolated single-SNP hits that showed significant p-values in both Discovery and Replication Cohorts. This approach minimises single-occurrence false positives that are caused by imputation or genotyping errors, whilst maintaining sensitivity to detect genuine associations that may be represented by a single variant.

## 3 Results

To identify latent subgroup structure in functional brain organisation, we first derived individual-specific resting-state networks (RSN) maps from 19,993 UK Biobank participants using stochastic PROFUMO (sPROFUMO). We then characterised continuous axes of spatial variation (“fingerprint features”) and identified latent subgroups from these features using unsupervised modelling. We next evaluated subgroup reproducibility, phenotypic relevance, patterns of subgroup variability in the spatial organisation of cognitive and sensory-motor RSNs and genetic relevance of subgroups, i.e., identifying links between patterns of subgroup differences and genetic variability in the spatial organisation of the RSNs.

### 3.1 Characterising subgroups based on the spatial topographies of the RSNs

We estimated 50 population-informed individualised RSNs by applying sPROFUMO to rfMRI data from 19,993 UKB subjects. As shown in **Figure 1**, each RSN’s spatial map was concatenated across subjects, yielding k = 50 matrices of size Nsubject by Nvoxel. These were then separately fed into two feature extraction algorithms: FMRIB’s Linked ICA (FLICA) (Gong et al., 2021) and Subject ICA. Whilst FLICA finds spatially-independent components (having spatially non-Gaussian distributions), Subject ICA finds subject-independent components (i.e., being non-Gaussian in the other dimension – subject-wise, not spatially).

FLICA was originally proposed to find Independent Components based on shared patterns of spatial variability across multiple imaging modalities. Here we adopted this technique to instead find shared patterns of variability across multiple RSNs. The resulting outputs are spatially independent components, which yield a parsimonious set of subject-specific fingerprints based on millions of features across voxels and RSNs.

We estimated 500 FLICA fingerprint features using the scalable bigFLICA implementation (2.2.2). As a preprocessing step, as elaborated in section 2.2, bigFLICA uses a linked PCA approach and Dictionary Learning to reduce data dimensionalities across subjects and voxels, respectively, before running the linked ICA. FLICA features are spatially independent; we next fed the FLICA outputs into “subject” ICA. Subject ICA uses subject by subject covariation patterns, and thus explicitly places subjects who are similar to each other closer to each other across the axes of variability – this approach can be expected to provide a particularly useful basis for subgroup modelling. We estimated 500 subjectICA fingerprint features. The number of fingerprint features was selected to match our previous studies (Farahibozorg et al., 2025, 2021). Together, these two complementary fingerprint representations capture both shared patterns of spatial variation across functional networks (FLICA) and similarities between individuals (subjectICA), providing 1,000 fingerprint features for subsequent latent subgroup discovery.

As shown in **Figure 1**, we then used each of these FLICA and subjectICA features as input to independent 1D Gaussian Mixture Models (GMMs), as elaborated in Methods section 2.2.3, to characterise subgroups. This approach identifies latent subgroup structure wherever the population distribution departs from Gaussianity. For example, if a feature’s distribution over subjects has a long tail on one or both sides, the subjects at each tail can be expected to be different from the main body of the distribution. Such subjects could represent smaller distinct subgroups.

As shown in **Figure 2**a,b, we used two metrics, Kurtosis and Skewness, to characterise non-Gaussianities of all the 500+500 fingerprint features. While 440/500 FLICA and 499/500 subjectICA components showed significant non-Gaussianity according to Kurtosis scores (after correcting p-values for multiple comparisons), only 83/500 FLICA and 32/500 subjectICA components (i.e., a minority) showed significant Skewness scores (**Figure 2**a,b). This indicates that a majority of the fingerprint features have approximately symmetrical distributions, with non-Gaussianity corresponding instead to super- (longer tails on both sides) or sub- (shorter tails on both sides) Gaussian behaviour. Therefore, deviations from Gaussianity were commonly found across the fingerprint features but were predominantly driven by changes in tails rather than asymmetry.

**Figure 2.**
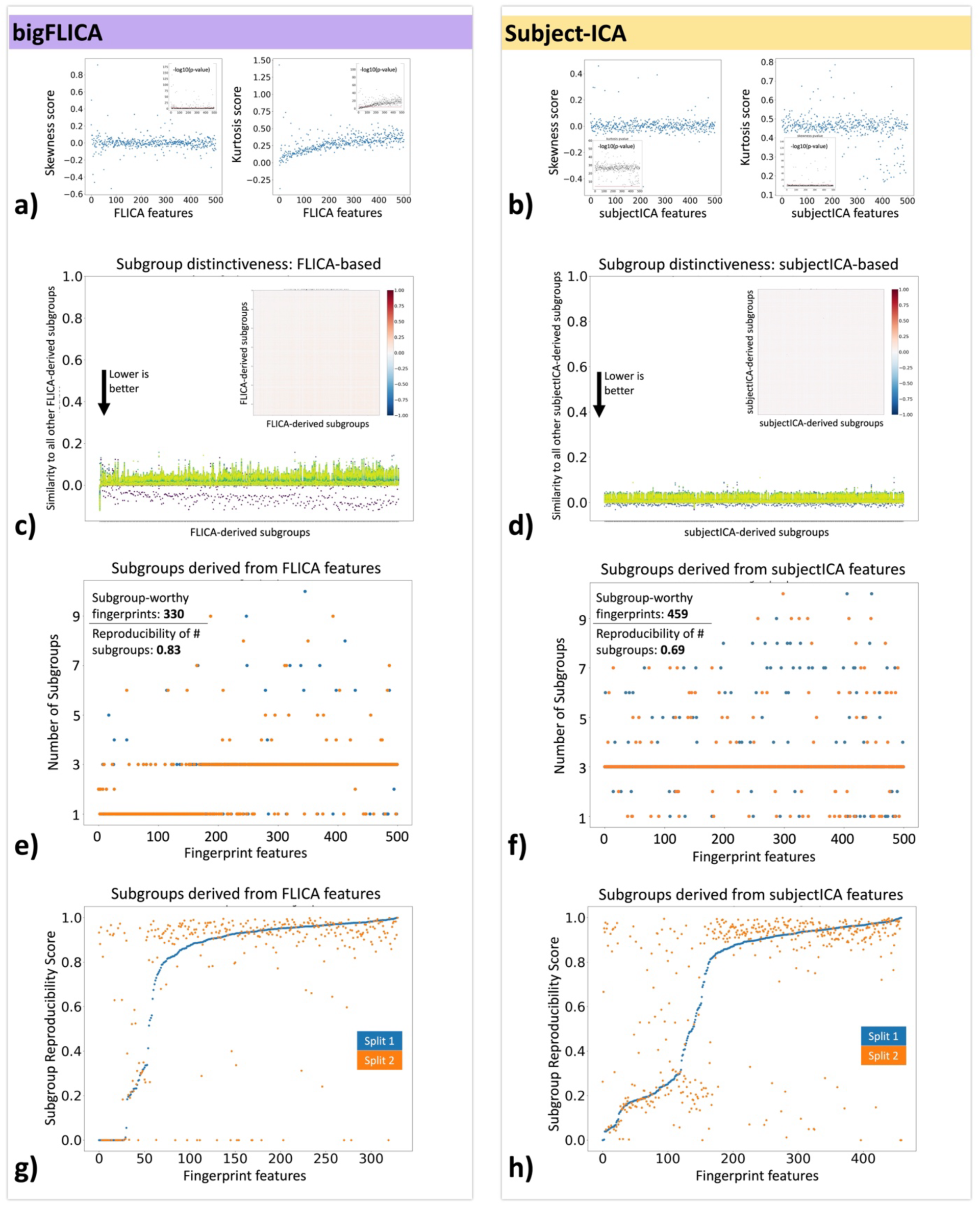
Validating the identified subgroups, their uniqueness and reproducibility. Left panel shows FLICA-based subgroups and right panel shows subjectICA-based subgroups. a, b) Characterising non-Gaussianities of the FLICA and subjectICA components (i.e., fingerprint features) using Skewness and Kurtosis scores (shown with blue datapoints) and respective p-values (black datapoints). A majority of the components (440 FLICA and 499 subjectICA) showed significant non-Gaussianity according to Kurtosis scores but only a minority of the components (83 FLICA and 32 subjectICA) showed significant non-Gaussianity according to Skewness scores. c, d) Using adjusted Rand index to compute how similar the subgroup definitions were across fingerprint features. Low scores denote highly distinct subgroup definitions, which is desired. Only features that identified more than 1 subgroup were included in this analysis. e, f) Split-half reproducibility of the number of subgroups, comparing split 1 (blue) and split 2 (orange) with full data. 330/500 FLICA and 459/500 subjectICA features were found to be subgroup-worthy, i.e., identifying more than 1 group. Reproducibility scores for the number of identified subgroups were found to be 0.83 and 0.69 for FLICA and subjectICA, respectively. g, h) Split-half reproducibility of subgroup definitions, comparing data splits with full data. Here we tested the similarity of subgroup definitions (i.e., subjects being assigned to the same subgroups) using adjusted Rand index. Each method yielded between 250 and 300 highly reproducible subgroup definitions.

After estimating the number of subgroups per fingerprint feature using 1D GMMs, we tested whether or not those subgroups were distinct across the features. This is shown in **Figure 2**c,d. We computed adjusted Rand index between subgroupings originating from each feature and subgroupings of every other feature – this metric computes similarities between a pair of subgroup definitions. Therefore, smaller values are desirable as they would indicate distinct subgroup definitions. The absolute values of the adjusted Rand indices were found to be 0.033±0.017 (max 0.071±0.03) for FLICA-based and 0.015±0.012 (max 0.04±0.066) for subjectICA-based subgroups, thus indicating highly distinct subgroups derived from various fingerprint features. This indicates that various subgroupings are capturing patterns beyond trivial and/or repetitive linear trends in the data.

To determine whether the identified subgroup structure reflected robust population organisation, we next examined split-half reproducibility of the subgroups. First, we tested the reproducibility of the number of subgroups. By dividing the 19,993 subjects into two halves and re-running the GMM separately for each half, we checked how many of the fingerprint features reproduced the same number of subgroups. This is shown in **Figure 2**e,f. Of the FLICA features, 330 identified > 1 subgroup in the full sample size. The number of subgroups per fingerprint feature were then computed independently for each split, and the reproducibility of the number of subgroups was found to be 0.83. Of the 500 subjectICA features, 459 identified > 1 subgroup in the full sample size, and reproducibility of the number of subgroups compared with data splits was 0.69.

We finally tested if the subjects that were assigned to the same subgroups in full data were also assigned to the same subgroups in split-halves. This was evaluated using adjusted Rand index, and results are shown in **Figure 2**g,h. Of the 330 subgroup definitions extracted from FLICA, 257 in data split 1 and 257 in data split 2 showed reproducibility scores higher than 0.8, and the overall reproducibility scores were 0.82±0.32 for both data splits. Of the 459 subgroup definitions extracted from subjectICA, 295 in data split 1 and 302 in data split 2 showed reproducibility scores higher than 0.8 and the overall reproducibility scores were 0.696±0.33 and 0.70±0.35 for splits 1 and 2, respectively.

Together, these analyses demonstrate that latent subgroup structure is common across functional fingerprint features and can be estimated reproducibly across subsets of the population. Whilst subjectICA identified a larger number of subgroup-relevant features, FLICA-derived subgroup definitions were generally more reproducible. Both approaches yielded approximately 250-300 highly reproducible subgroup structures, providing a robust foundation for subsequent phenotypic, spatial and genetic analyses.

### 3.2 Phenotypical relevance of subgroups

#### 3.2.1 Phenotypic differences between subgroups

As elaborated earlier, fingerprint features and subgroups were identified using unsupervised methods, and thus are completely uninformed of subjects’ cognition, lifestyle, physical or mental health. We next asked whether the latent subgroups identified purely from functional brain organisation also captured meaningful variation in non-imaging phenotypes. For this purpose, we conducted statistical comparisons between subgroups with respect to the following non-imaging derived phenotype (nIDP) categories: cognition (68 nIDPs), mental health (142 nIDPs), alcohol (37 nIDPs) and tobacco (19 nIDPs) consumption scores, cardiovascular (77 nIDPs) and bone (93 nIDPs) health scores. Results are shown in **Figure 3**a. As illustrated earlier, each FLICA/subjectICA fingerprint feature yielded Kj subgroups. We compared each nIDP (e.g., fluid intelligence) between these subgroups using Kruskal-Wallis test and computed the corresponding p-values.

**Figure 3.**
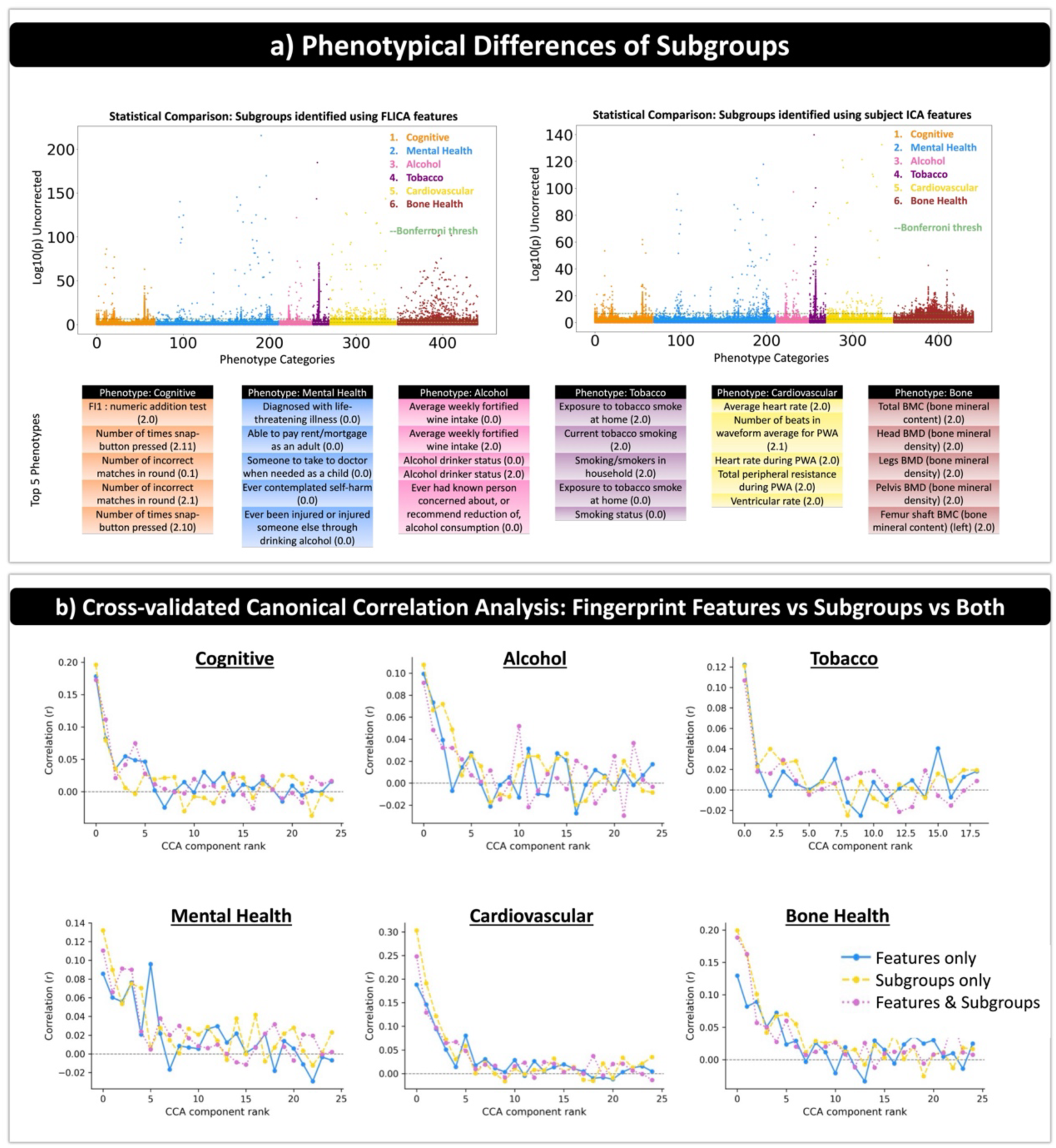
Phenotypical relevance of subgroups. a) Statistical comparisons of phenotypes were conducted between various subgroup definitions and -log10(p-values) are shown for FLICA (left) and subjectICA (right). Dashed green lines show Bonferroni threshold. The top 5 phenotypes within each category (highest number of smallest p-values) are shown in the tables. b) Canonical Correlation Analysis (CCA) with 5-fold cross-validation was performed to find axes of co-variation between brain-based features and phenotypes. Specifically, we compared brain-based features defined as features-only, subgroups-only and subgroups+features. For 5 out of 6 phenotype categories – cognitive, alcohol, cardiovascular, bone and mental health – subgroups-only provided the top-matched CCA component. For the tobacco category, features-only and subgroups-only provided the top-matched CCA component and were on par. For visualisation purposes, only up to 25 CCA components are illustrated on the x-axis.

The -log(10) of the p-values is illustrated in **Figure 3**a, with the Bonferroni threshold (correcting across features x nIDPs) shown as the dashed green line – any data points above that threshold are considered significant. As can be noted from these graphs, both FLICA-based and subjectICA-based subgroupings showed hundreds of significant nIDP-hits across the six categories. **nIDP-hit**s refer to the significant nIDP differences between subgroups derived from a single fingerprint feature, after correction for multiple comparisons, see 2.2.6 for details.

Related to cognition and mental health: first, within the cognition category, we found 877 FLICA-related and 487 subjectICA-related nIDP-hits. Some of the strongest nIDP-hits were related to fluid intelligence, reaction time and working memory performance. Second, within the mental health category, we found 522 FLICA-related and 293 subjectICA-related nIDP-hits. Some of the top nIDP-hits were related to life-threatening disease diagnosis, ability to pay rent/mortgage, someone to take to the doctor as a child, ever contemplating self-harm, and ever injured or being injured due to alcohol. Interestingly, several of the most significant nIDP-hits under this category can be linked to socioeconomic background.

Related to lifestyle and substance use: first, within the alcohol category, we found 314 FLICA-related and 214 subjectICA-related nIDP-hits. Some of the top nIDP-hits were related to alcohol drinking status, concerns about level of alcohol consumption, and fortified wine intake. Second, within the tobacco category, we found 587 FLICA-related and 591 subjectICA-related nIDP-hits. The top nIDP-hits were related to current smoking status and exposure to tobacco smoke at home.

Related to physical health: first, within the cardiovascular health category, we found 280 FLICA-related and 127 subjectICA-related nIDP-hits. The top 5 nIDP-hits were related to heart rate, number of beats during Pulse Wave Analysis and ventricular rate. Second, within the bone health category, we found 487 FLICA-related and 903 subjectICA-related nIDP-hits. The top 5 nIDP-hits were related to Bone mineral density and content in various body parts.

Put together, out of 436×459 (nIDP x Nfeature) comparisons for subjectICA, 2,615 p-values and out of 436×330 (nIDP x Nfeature) comparisons for FLICA, 3,067 p-values passed the Bonferroni threshold. Therefore, despite subgroup discovery being performed entirely without phenotype information, approximately 5,700 significant subgroup-phenotype associations were identified across cognition, lifestyle, physical and mental health. These findings demonstrate that latent subgroup structure derived solely from functional brain organisation captures broad and biolosically relevant population heterogeneity rather than variation confined to a single phenotype domain.

#### 3.2.2 Canonical Correlation Analysis

We next used Canonical Correlation Analysis (CCA) to compare multivariate associations between each of the six phenotype categories and: a) subgroups; b) continuous features; or c) both representations concatenated. The six phenotype categories were the same as above. The goal here was to compare the shared variance between phenotypes and subgroup estimations, versus continuous fingerprint features, versus both combined. This was done to test whether the new subgroup modelling used in this study provides links to the non-imaging traits, above and beyond what can be achieved with the more commonly-used continuous features. CCA finds axes of population co-variations (canonical variates) between the brain-based features and phenotypes. This is done through linear projections of the two sets, such that the similarity between the transformed outputs is maximised (Methods section 2.2.8). To avoid overfitting and obtain generalisable results, we conducted cross-validated CCA with 5-fold cross-validation. We estimated shared variances for each pairwise comparison by computing correlations between the corresponding canonical covariates within the held-out test sets, and averaging those correlations across folds. These results are shown in **Figure 3**b.

The correlation values for the top-matched CCA components of the features-only CCA, subgroups-only CCA and both concatenated CCA, respectively, were as follows: 1) cognitive nIDPs: 0.178, 0.196 and 0.172; 2) mental health nIDPs: 0.096, 0.132 and 0.111; 3) alcohol nIDPs: 0.099, 0.108 and 0.091; 4) tobacco nIDPs: 0.122, 0.121 and 0.107; 5) cardiovascular nIDPs: 0.188, 0.303 and 0.248; 6) bone health nIDPs: 0.130, 0.199 and 0.189.

Therefore, for five of the six phenotype categories – cognitive, alcohol, cardiovascular, bone and mental health – subgroup modelling provided stronger brain-phenotype associations than continuous fingerprint features alone and combining the two did not further improve performance. This finding suggests that the discrete subgroup representation captures behaviourally relevant population structure more effectively than continuous fingerprints, and potentially reduces noise and redundant variability.

### 3.3 Spatial Maps of Subgroup Differences (MSDs) in the brain

To investigate the biolosical organisation of the discovered subgroups, we next examined how the canonical resting state networks (RSNs) varied spatially across the discovered subgroups. This was investigated from two angles, first to test if Maps of Subgroup Differences (MSDs) were reflective of the brain networks’ meta functionalities (i.e., sensorimotor vs higher-level cognitive processing). Second, we tested whether the MSDs reflected phenotypical differences; e.g., if certain phenotype categories showed similar patterns of subgroup differences across the brain voxels.

#### 3.3.1 MSDs: sensorimotor vs higher-level cognitive networks

As shown in **Figure 4**a, we examined MSDs in 11 canonical RSNs, consisting of 6 cognitive and 5 sensory-motor RSNs: cognitive networks included Default Mode 1&2, left and right Fronto-parietal, Ventral Attention and Language, sensory-motor networks included Visual 1&2, Motor 1&2 and Auditory. These RSNs were manually selected as some of the most well-known RSNs to represent cognitive and sensorimotor networks, respectively (Kong et al., 2025). For each RSN, we used subject-specific spatial maps, as estimated using sPROFUMO, and conducted voxel-wise statistical comparisons between the Kj subgroups identified using each of the j fingerprints. To keep the computational costs manageable, we restricted this analysis to the nIDP-hits (i.e., significant nIDP differences between subgroups derived from a single fingerprint feature, after correction for multiple comparisons), see **Figure 3**a, and section 2.2.7 for details. MSDs were obtained by: first, computing maps of significant differences between subgroups derived from each feature after correction for multiple comparisons; next, finding brain areas in which significant differences were most frequently observed across subgroups derived from various features; finally, binarising these maps at the significance threshold and accumulating the results across all the fingerprint features, as shown in **Figure 4**a left.

**Figure 4.**
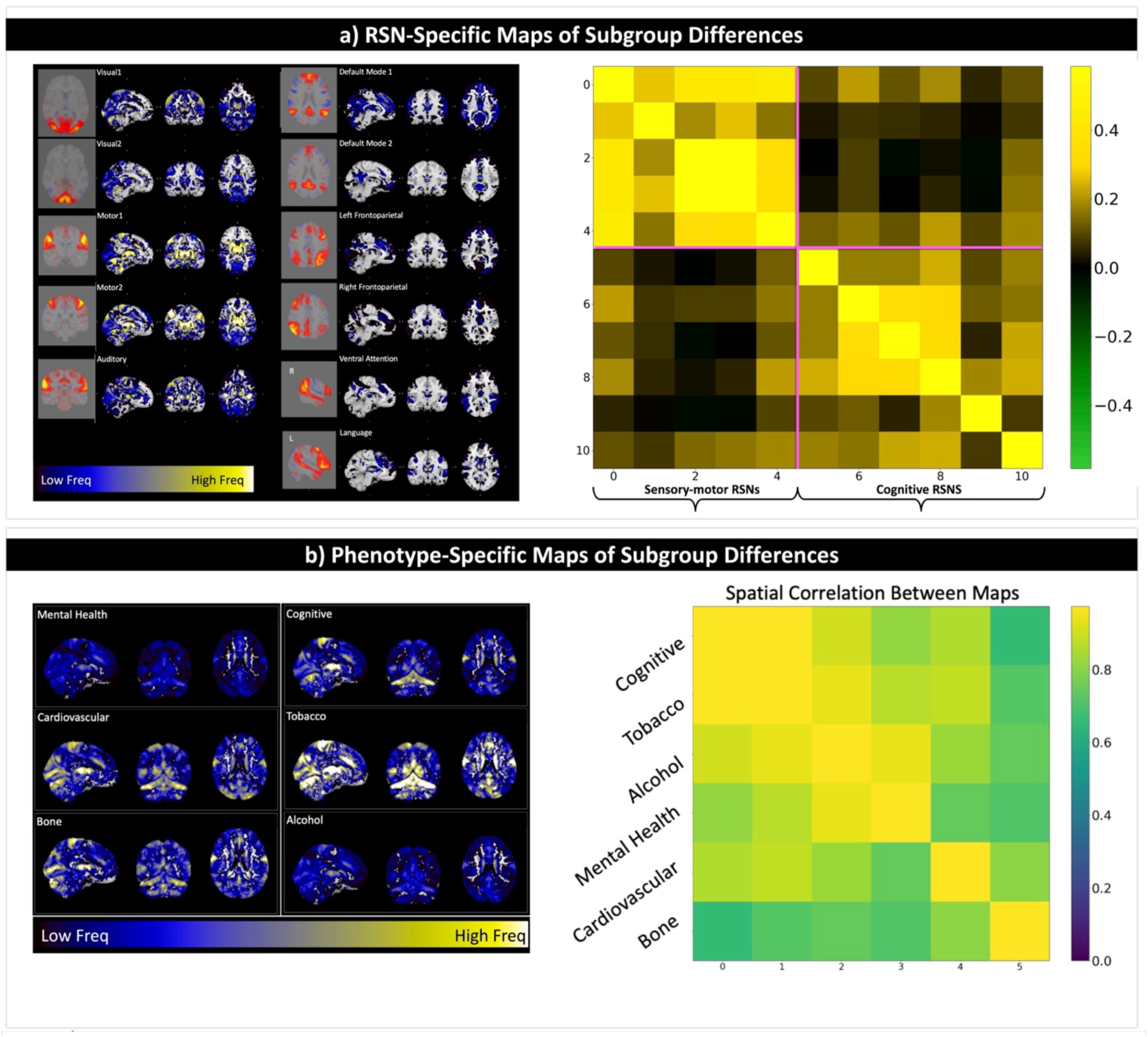
Spatial Maps of Subgroup Differences (MSD) in the brain. a) RSN-specific MSDs: Individual-specific RSNs estimated using sPROFUMO (i.e., PFMs) were used to conduct voxel-wise statistical comparisons between the subgroups, in order to localise the key areas of subgroup variability in the brain. Spatial maps of p-values were generated for subgroups derived from each fingerprint feature, binarised after correction for multiple comparisons, and accumulated across fingerprint features. The left panel shows MSDs estimated for 11 RSNs, and the right panel shows the spatial correlations between these MSDs. An interesting block structure was found whereby sensory-motor MSDs were similar to each other and distinct from MSDs derived from cognitive RSNs, and vice versa. b) Phenotype-specific MSDs: similar to panel (a), binarised maps of subgroup differences accumulated per phenotype category and across all the 11 RSNs. The left panel shows MSDs estimated for the 6 phenotype categories, and the right panel shows the spatial correlations between these MSDs. Cognitive- and tobacco-specific MSDs were highly similar to each other, alcohol- and mental health-specific MSDs were highly similar to each other, and cardiovascular-and bone-specific MSDs were relatively distinct from the other five categories.

MSDs showed a clear block structure separating sensory-motor and higher-order cognitive networks (**Figure 4**a right). Specifically, within-sensory correlations were found to be 0.35±0.12, within-cognitive were 0.19±0.08 and between sensory and cognitive were 0.08±0.07. Therefore, MSDs of the sensory-motor networks (Visual, Auditory and Motor) were similar to each other and distinct from MSDs of the cognitive networks, and vice versa. This is particularly notable because sensory-motor networks cover very different brain areas and are largely non-overlapping. Despite this, brain areas affected by subgroup differences in these networks were found to be similar to each other. Brain regions that showed the highest MSD values included subcortex, cerebellum, parieto-occipital and central regions. This was also observed for cognitive networks, whereby despite the networks being largely non-overlapping, the areas impacted by subgroup differences showed similarities across these networks and predominately covered frontal lobe, lateral and medial parietal lobe.

#### 3.3.2 MSDs: contrasting the six phenotype categories

Next, we examined the extent to which MSDs were reflective of phenotypical differences; i.e., computing and contrasting MSDs separately for the six phenotype categories. MSDs were computed using a similar process as above, with the exception that the binarised maps of significant differences were accumulated across nIDP-hits and RSNs *per phenotype category,* thus yielding MSDs that were specific to each phenotype category.

As shown in **Figure 4**b, here we found that cognitive and tobacco related spatial maps were very similar to each other (r = 0.97) and more distinct from the other four categories (r = 0.83±0.09). Medial frontal, occipital, central, cerebellar and subcortical brain regions were most pronounced in these maps. Additionally, mental health and alcohol related MSDs were highly similar to each other (r = 0.94) and relatively distinct from the other four maps (r = 0.82±0.076). These MSDs appeared to be more uniformly distributed across the brain regions. MSDs related to cardiovascular (r = 0.83±0.048) and bone health (r = 0.73±0.047) were relatively distinct from the other five categories. Together, these findings indicate that different categories of behavioural and health phenotypes are associated with distinct spatial patterns of subgroup variability rather than a common spatial configuration.

### 3.4 Genetic relevance of fingerprints and subgroups

Having established the link between subgroups and the individualised spatial organisation of brain networks, as well as their link to phenotypes, we next examined the relationship between subgroups and genetics. Demonstrating such correspondence would provide an independent biolosical validation of the latent subgroup structure identified using unsupervised methods from fMRI. To address this, we first identified genetic variants associated with the continuous fingerprint features (using Genome Wide Association Studies (GWAS)) and then tested whether spatial patterns of subgroup variability corresponded to regional patterns of genetic variation across functional brain networks. The latter was done by computing RSN-specific Maps of Allele Dosage (MAD) for SNP-hits in GWAS, and comparing those to MSDs from the previous analyses.

#### 3.4.1 **Summary of** GWAS **results**

GWAS requires continuous features as input, and therefore it was performed per fingerprint feature, whereby univariate associations between each fingerprint feature and over 90 million imputed genetic variants were computed. Results were corrected for multiple comparisons. Data were deconfounded using both imaging and genetics confounds, as described in Methods. Of the 19,993 subjects included in this study, ∼16,000 were included in UKB’s BIG40 (updated with 63k subjects in 2025) sample, after applying genetic and ancestry filters. We divided the sample into Discovery (n = 11,198) and Replication (n = 5,583) cohorts. **Figure 5**a shows the Manhattan plot of GWAS for one example FLICA fingerprint feature; the same analysis was repeated across all the 500+500 FLICA and subjectICA fingerprint features. Based on the standard GWAS threshold (−log10 (p-value) = 7.5) and after prefiltering as outlined in 2.2.9, we identified 58 (38 FLICA-based and 20 subjectICA-based) significant Single Nucleotide Polymorphism (SNP) hits, 13 of which were replicated in the replication cohort.

**Figure 5.**
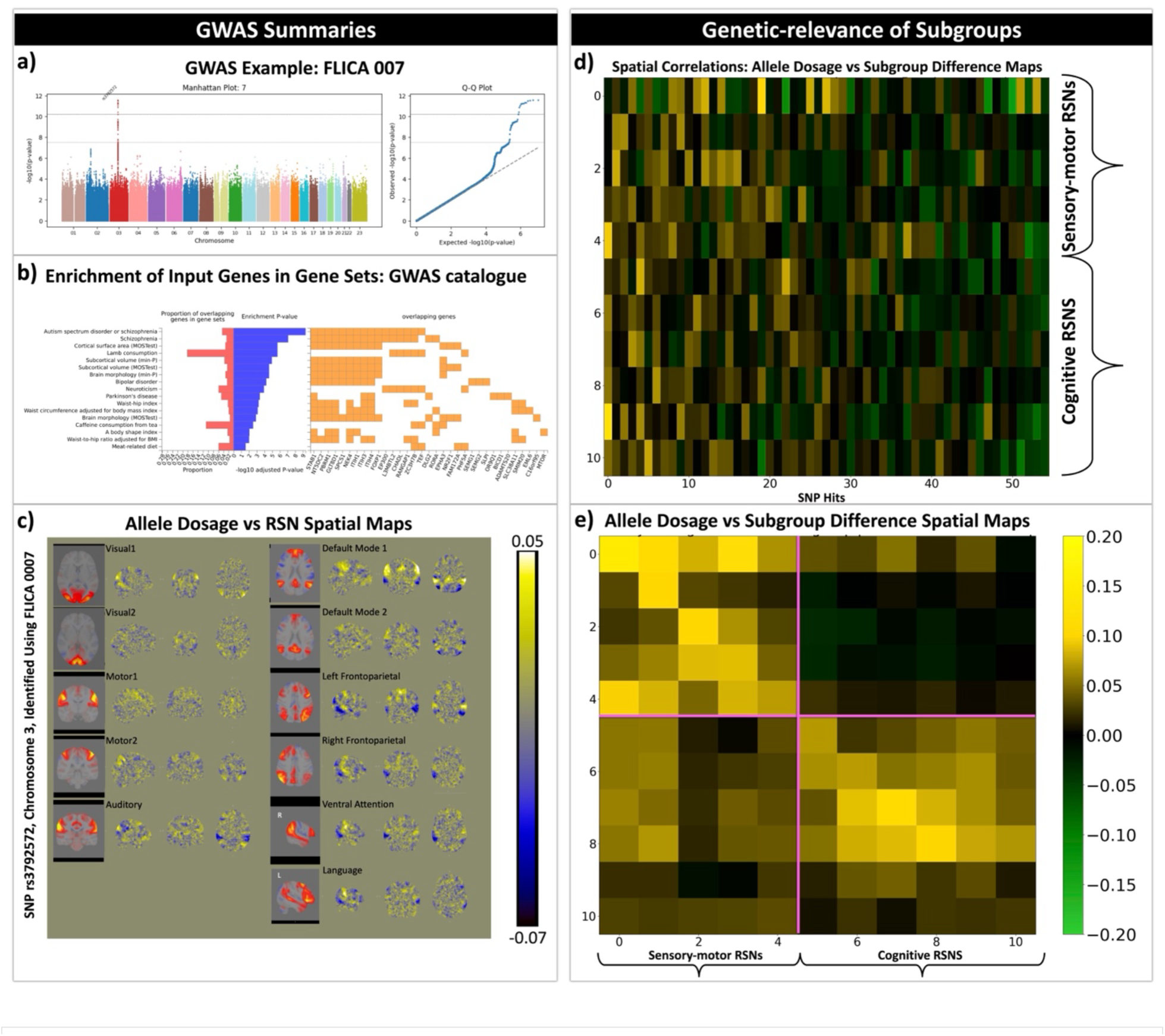
Genetic relevance of fingerprints and subgroups: a) A Manhattan plot of a GWAS conducted on an example fingerprint feature (FLICA component 007). The same analysis was conducted across all FLICA and subjectICA fingerprint features, and significant clusters of SNP-hits were identified (after correction for multiple comparisons). b) FUMA software was used to link the SNP-hits to Genes and Genes to functions and disease. Based on the GWAS catalogue that is used as reference in FUMA software, we found significant enrichment of the identified genes to gene sets that have been previously linked to brain-related disorders such as Autism Spectrum Disorder and Schizophrenia, brain anatomy and lifestyle related to diet and body shape. c) Spatial Maps of Allele Dosage (MAD) were calculated as voxel-wise correlations between subject-specific RSNs and subject-specific allele dosage for each SNP-hit. Results are shown for 11 RSNs of interest for an example SNP hit. d) RSN-specific MADs were spatially correlated with their corresponding Maps of Subgroup Differences (MSDs, **Figure 4**a), results per SNP-hit and RSN are shown. e) MADs were averaged across SNP-hits per RSN and spatially correlated with the corresponding MSDs. Diagonal values were found to be 0.103±0.026 for sensory-motor RSNs and 0.069±0.03 for cognitive RSNs.

We subsequently used FUMA (Watanabe et al., 2017) to link the SNPs to genes, and the genes to functions and disease. The SNP2GENE analysis identified 54 genomic risk loci and 69 mapped genes. Of the SNPs, 51.3% were intergenic and showed significant enrichment (enrichment value = 1.10, p = 2.18 × 10⁻⁷), while 32.2% were intronic and showed significant depletion (enrichment value = 0.885, p = 1.73 × 10⁻⁶). The remaining SNPs did not show significant functional annotation enrichment. Subsequent GENE2FUNC analysis compared the identified genes with those reported in GWAS catalogue and found 17 significant overlaps with gene sets (**Figure 5**b). These gene sets were related to three categories of function/diseases: a) brain-related disorders and traits, including Autism spectrum disorder, Schizophrenia, Bipolar disorder, Neuroticism and Parkinson’s; b) brain anatomy, including cortical surface area, subcortical volume and brain morphology; c) diet and body shape, including meat and caffein consumption, waist-hip index and body shape index.

#### 3.4.2 RSN-specific spatial maps of genetics versus subgroup variability

To determine whether subgroup variability and genetic variation showed similar spatial organisation, we next estimated spatial Maps of Allele Dosage (MADs) for each significant SNP and compared them with the corresponding Maps of Subgroup Differences (MSDs) found in 3.3. Specifically, subject-specific Allele dosage of SNP-hit matrices (Nsubject x NSNP_hit) were correlated with subject-specific PFM spatial maps (Nsubject x Nvoxel matrices per RSN) to obtain NSNP_hit voxel-wise MADs. An example set of maps corresponding to one SNP-hit is shown in **Figure 5**c. For each RSN, we then found the spatial correlations between MADs with corresponding MSDs (**Figure 4**a-left), the results of which are shown in **Figure 5**d.

Finally, **Figure 5**e shows the correlations between RSN-specific MADs (averaged across SNP-hits) and MSDs. The diagonal of this correlation matrix captures the one-to-one correspondence between the allele dosage and subgroup difference maps and were found to be: 0.103±0.026 for sensory-motor RSNs and 0.069±0.03 for cognitive RSNs. Consistent with the MSD analyses (Section 3.3), here as well we found a block structure, with stronger similarities within sensory-motor networks (r = 0.074±0.038) and within higher-order cognitive networks (r = 0.057±0.025) than between these functional systems (r = 0.001±0.021).

Together, these analyses demonstrate that the functional fingerprint features capture genetic variation associated with brain-related traits and disorders, while the spatial organisation of subgroup differences corresponds to regional patterns of genetic variability across canonical functional networks. These findings provide independent biolosical support for the latent subgroup structure identified from functional neuroimaging, and suggest that this organisation reflects meaningful axes of population variability.

## 4 Discussion

Using individualised functional brain topographies from 19,993 UK Biobank participants, we demonstrate that large-scale functional neuroimaging contains reproducible latent subgroup structure that bridges population-average and individual-specific models of the brain’s functional organisation. To reveal this, we developed a scalable framework that builds on stochastic Probabilistic Functional Modes (sPROFUMO) (Farahibozorg et al., 2021), combined with two unsupervised fingerprint extraction approaches (bigFLICA and subjectICA) and feature-wise (1D) Gaussian Mixture Modelling to identify multiple subgroup definitions across distinct axes of functional variation. Importantly, unlike conventional approaches based on population-average models or temporal functional connectivity, our framework uses *individualised spatial topographies* of functional brain networks to characterise latent population structure. Across 19,993 participants, we identified 1,000 fingerprint features and hundreds of reproducible subgroup definitions that were associated with ∼5,700 significant phenotype differences, exhibited distinct organisation across sensory-motor and higher-order cognitive systems, and showed correspondence with regional patterns of genetic variation across the brain.

Subgroup modelling based on large-scale population imaging can take a significant step towards a core aim of fMRI research: providing biomarkers of traits and disease while helping bridge the gap between population neuroimaging and precision medicine (Demeter and Greene, 2025; Parkes et al., 2020). Specifically, conventional neuroimaging datasets with small sample sizes typically included homogeneous cohorts (Button et al., 2013; Szucs and Ioannidis, 2020). As a result, conventional fMRI modelling techniques were designed to analyse such datasets, and therefore relied on population average models to identify consensus patterns within groups (Calhoun et al., 2001; Yeo et al., 2011), rather than the structured heterogeneity that becomes apparent in large-scale datasets.

However, compelling evidence accumulated over recent years has established structured and meaningful patterns of individual variability in brain function and connectivity (Gordon et al., 2017; Kong et al., 2019). This has led to extensive research into fMRI-based fingerprinting (Finn et al., 2015). However, to fully exploit the potential of large heterogeneous populations in modelling various sources of subject variability, it is important to bridge the gap between population-average and individual-specific modelling – this can be achieved using stratified methods. Subgroup modelling, as proposed in the current paper, takes a step towards bridging this gap. Specifically, using multiple phenotype categories – cognition, mental health, alcohol and tobacco use, bone and cardiovascular health – we identified thousands of highly significant differences between the data-driven subgroups. This provides a proof-of-principle that incorporating latent subgroup structure into models of brain function captures biolosically meaningful population heterogeneity and may provide a richer foundation for future individualised fMRI biomarkers.

Subgroup modelling has been used in previous fMRI literature for applications such as brain disorder classification (Du et al., 2018) and disease subtyping (Brucar et al., 2023; Miranda et al., 2021). However, several challenges hinder the generalisability and reproducibility of such approaches, including limited sample sizes or the lower quality of clinical datasets. Large-scale high-quality datasets of thousands of healthy participants will, by definition, include hundreds of potential clinical cohorts in pre-diagnosis stages. For example, by 2027, in the imaged cohort of the UK Biobank, 6,000, 4,000 and 2,800 participants are expected to develop Alzheimer’s disease, stroke and Parkinson’s, respectively (Miller et al., 2016). Therefore, these datasets are not only useful for characterising healthy populations at scale but also provide a solid foundation for modelling clinical data. Specifically, for Bayesian methods such as sPROFUMO, UK Biobank can provide informative “priors” for subsequent disease classification and subtyping, whereby the state-of-the-art transfer learning methods can be used to inform disease classification and subtyping in any new clinical data, based on similarities to the subpopulations that were identified in the current study.

The approach presented here is based on individualised spatial topographies of the brain’s functional networks. Compared with functional connectivity, spatial maps remain a much less explored dimension of individual variability (due to the limitations of the conventional methods), but recent studies have revealed crucial aspects of subject variability being captured by spatial maps (Bijsterbosch et al., 2018; Farahibozorg et al., 2025; Kong et al., 2019). Functional connectivity analysis computes temporal correlations between the brain networks (or parcels), and as such, it cannot be used to “localise” the network anomalies. This hinders biolosical interpretability of the functional connectivity-based biomarkers. In contrast, in this study we used fully individualised spatial maps to obtain voxel-wise maps of subgroup differences (MSD), as shown in **Figure 4**. Interestingly, we found distinct MSDs for sensory-motor versus cognitive networks, whereby MSDs related to visual, auditory and motor networks were highly similar to each other, and distinct from MSDs related to the default mode, fronto-parietal, ventral attention and language networks, and vice versa. This block structure in the spatial correlations between MSDs suggests that subgroup variability is organised at the level of meta-networks rather than individual functional networks.

This meta-network organisation aligns with previous investigations into amplitudes of the brain networks (Bijsterbosch et al., 2017) as well as in the dynamic states (referred to as meta-states in (Vidaurre et al., 2017)). This finding can be interpreted in the context of the growing evidence that inter-individual variability in functional brain organisation is anatomically-informed (rather than random or uniform), and thus differs between sensory-motor and associative systems (Finn et al., 2015; Mueller et al., 2013). For sensory-motor networks, the areas that we found in this study, including subcortical, cerebellar, and parieto-occipital regions, reflect the relatively hard-wired developmental trajectories and structural-functional connectivity that characterise the unimodal brain regions (Baum et al., 2020; Guell et al., 2018). In contrast, we found frontal and parietal association cortices highlighted in the MSDs related to the cognitive networks - consistent with these regions maturing later during development, showing higher synaptic plasticity, and greater inter-individual variability in functional connectivity (Mueller et al., 2013; Sydnor et al., 2021). This is broadly consistent with the cortical hierarchy framework, where transmodal association areas are thought to be less constrained by genetics and more sensitive to environmental factors (Margulies et al., 2016; Sydnor et al., 2021). Together, these results suggest that subgroup difference maps identified by our unsupervised techniques reflect biolosically meaningful axes of population variability, where the spatial organisation of these MSDs is broadly aligned with the well-known functional organisation of the cortex.

The genetic analysis provides independent evidence supporting the biolosical relevance of these subgroup differences. As well as identifying genetic loci associated with continuous fingerprint features, we found clear spatial correspondence between maps of subgroup differences and regional patterns of genetic variation across the same functional networks. Although these spatial associations do not imply causal mechanisms, they suggest that the spatial organisation of subgroup variability is partly aligned with the genetic architecture underlying functional brain organisation, complementing the behavioural and spatial evidence presented here.

Beyond characterising normative population variability, the subgroup organisation identified here may also have implications for understanding disease vulnerability. The prefrontal and parietal association cortex, subcortex, and cerebellum that were found here as the main zones of cross-subgroup differences, have all been implicated in various neuropsychiatric conditions (Goodkind et al., 2015; Sha et al., 2019), raising the possibility that the same pathways that accommodate subgroup diversity in healthy populations may also constitute points of vulnerability in diseases. More broadly, these findings provide a framework for investigating heterogeneity in both healthy populations and clinical cohorts using latent subgroup structure derived from individualised functional brain organisation.

## 5 Acknowledgements

## Author contribution

Conceptualisation, Investigation, Resources, Funding acquisition, Writing—review & editing: SRF, MWW, SMS; Methodology, Software: SRF, SMS, LTE; Validation, Data curation, Formal Analysis and Visualisation, Writing—original draft: SRF.

## Data availability

- UK Biobank data is available upon registration and applying for data access from UK Biobank website: http://www.ukbiobank.ac.uk/register-apply.

## Code availability

- Code for sPROFUMO is currently available from the following repository, it will be made available in an upcoming FSL release: https://git.fmrib.ox.ac.uk/profumo/profumo/-/tree/sprofumo-cpp-clean.

## Use of Generative AI

Some parts of the manuscript and analysis scripts were fine-tuned with the assistance of AI tools (e.g., ChatGPT Edu, OpenAI) to improve coding style, text clarity and language. Outputs were carefully reviewed and validated by the authors.

## 7 Supplementary Materials

**Table S 1.** Confounds.

| Confound # | Confound Name |
| --- | --- |
| 1 | 'Age_Site_1' |
| 2 | 'Age_Site_2' |
| 3 | 'Age_Site_3' |
| 4 | 'Sex_1_Site_1' |
| 5 | 'Sex_1_Site_2' |
| 6 | 'Sex_1_Site_3' |
| 7 | 'HeadSize_Site_1' |
| 8 | 'HeadSize_Site_2' |
| 9 | 'HeadSize_Site_3' |
| 10 | 'TablePos_COG_Z_Site_1' |
| 11 | 'TablePos_COG_Z_Site_2' |
| 12 | 'TablePos_COG_Z_Site_3' |
| 13 | 'D_clean_p90_Site_1' |
| 14 | 'D_clean_p90_Site_2' |
| 15 | 'D_clean_p90_Site_3' |
| 16 | 'Site_1_vs_2' |
| 17 | 'Site_1_vs_3' |
| 18 | 'T1_and_T2_for_FreeSurfer_1_Site_1' |
| 19 | 'SCALING_T1_1_Site_1' |
| 20 | 'SCALING_rfMRI_1_Site_1' |
| 21 | 'SCALING_tfMRI_1_Site_1' |
| 22 | 'S_clean_p90_Site_1' |
| 23 | 'SD_clean_mean_Site_1' |
| 24 | 'SD_clean_p90_Site_1' |
| 25 | 'TablePos_Table_Site_1' |

**Table S 2.** Cognitive nIDPs.

| IDP # | IDP Name | IDP # | IDP Name |
| --- | --- | --- | --- |
| 1 | Number of incorrect matches in round (0.1) | 35 | Time elapsed (2.6) |
| 2 | Number of incorrect matches in round (0.2) | 36 | Time elapsed (2.7) |
| 3 | Number of incorrect matches in round (2.1) | 37 | Time elapsed (2.8) |
| 4 | Number of incorrect matches in round (2.2) | 38 | Digits entered correctly (2.0) |
| 5 | Number of incorrect matches in round (2.3) | 39 | Digits entered correctly (2.1) |
| 6 | Number of times snap-button pressed (0.0) | 40 | Digits entered correctly (2.2) |
| 7 | Number of times snap-button pressed (0.1) | 41 | Digits entered correctly (2.3) |
| 8 | Number of times snap-button pressed (0.2) | 42 | Digits entered correctly (2.4) |
| 9 | Number of times snap-button pressed (0.3) | 43 | Digits entered correctly (2.5) |
| 10 | Number of times snap-button pressed (0.4) | 44 | Digits entered correctly (2.6) |
| 11 | Number of times snap-button pressed (0.10) | 45 | Digits entered correctly (2.7) |
| 12 | Number of times snap-button pressed (0.11) | 46 | Total errors traversing numeric path (trail #1) (2.0) |
| 13 | Number of times snap-button pressed (2.0) | 47 | Mean time to correctly identify matches (0.0) |
| 14 | Number of times snap-button pressed (2.1) | 48 | Fluid intelligence score (0.0) |
| 15 | Number of times snap-button pressed (2.2) | 49 | Fluid intelligence score (2.0) |
| 16 | Number of times snap-button pressed (2.3) | 50 | Duration to complete numeric path (trail #1) (0.0) |
| 17 | Number of times snap-button pressed (2.4) | 51 | Duration to complete alphanumeric path (trail #2) (0.0) |
| 18 | Number of times snap-button pressed (2.5) | 52 | Digits entered correctly (2.8) |
| 19 | Number of times snap-button pressed (2.7) | 53 | Maximum digits remembered correctly (2.0) |
| 20 | Number of times snap-button pressed (2.10) | 54 | Attempted fluid intelligence (FI) test. (2.0) |
| 21 | Number of times snap-button pressed (2.11) | 55 | FI1 : numeric addition test (0.0) |
| 22 | Prospective memory result (0.0) | 56 | FI1 : numeric addition test (2.0) |
| 23 | Prospective memory result (2.0) | 57 | FI3 : word interpolation (0.0) |
| 24 | FI8 : chained arithmetic (2.0) | 58 | FI3 : word interpolation (2.0) |
| 25 | FI9 : concept interpolation (2.0) | 59 | FI4 : positional arithmetic (0.0) |
| 26 | Duration to complete numeric path (trail #1) (2.0) | 60 | FI4 : positional arithmetic (2.0) |
| 27 | Mean time to correctly identify matches (2.0) | 61 | FI5 : family relationship calculation (0.0) |
| 28 | Number of incorrect matches in round (0.0) | 62 | FI5 : family relationship calculation (2.0) |
| 29 | Time elapsed (2.0) | 63 | FI6 : conditional arithmetic (0.0) |
| 30 | Time elapsed (2.1) | 64 | FI6 : conditional arithmetic (2.0) |
| 31 | Time elapsed (2.2) | 65 | FI7 : synonym (0.0) |
| 32 | Time elapsed (2.3) | 66 | FI7 : synonym (2.0) |
| 33 | Time elapsed (2.4) | 67 | Duration to complete alphanumeric path (trail #2) (2.0) |
| 34 | Time elapsed (2.5) | 68 | Number of puzzles correctly solved (2.0) |

**Table S 3.** Mental health nIDPs.

| IDP # | IDP Name | IDP # | IDP Name |
| --- | --- | --- | --- |
| 1 | 'Mood swings (0.0)', | 72 | 'Difficulty concentrating during worst depression (0.0)', |
| 2 | 'Mood swings (2.0)', | 73 | 'Fraction of day affected during worst episode of depression (0.0)', |
| 3 | 'Miserableness (0.0)', | 74 | 'Thoughts of death during worst depression (0.0)', |
| 4 | 'Miserableness (2.0)', | 75 | 'Duration of worst depression (0.0)', |
| 5 | 'Irritability (0.0)', | 76 | 'Frequency of depressed days during worst episode of depression (0.0)', |
| 6 | 'Irritability (2.0)', | 77 | 'Impact on normal roles during worst period of depression (0.0)', |
| 7 | 'Sensitivity / hurt feelings (0.0)', | 78 | 'Ever had prolonged loss of interest in normal activities (0.0)', |
| 8 | 'Sensitivity / hurt feelings (2.0)', | 79 | 'Lifetime number of depressed periods (0.0)', |
| 9 | 'Fed-up feelings (0.0)', | 80 | 'Depression possibly related to childbirth (0.0)', |
| 10 | 'Fed-up feelings (2.0)', | 81 | 'Ever had prolonged feelings of sadness or depression (0.0)', |
| 11 | 'Nervous feelings (0.0)', | 82 | 'Depression possibly related to stressful or traumatic event (0.0)', |
| 12 | 'Nervous feelings (2.0)', | 83 | 'Professional informed about depression (0.0)', |
| 13 | 'Worrier / anxious feelings (0.0)', | 84 | 'Feelings of tiredness during worst episode of depression (0.0)', |
| 14 | 'Worrier / anxious feelings (2.0)', | 85 | 'Feelings of worthlessness during worst period of depression (0.0)', |
| 15 | 'Tense / 'highly strung' (0.0)', | 86 | 'Ever taken cannabis (0.0)', |
| 16 | 'Tense / 'highly strung' (2.0)', | 87 | 'General happiness (0.0)', |
| 17 | 'Worry too long after embarrassment (0.0)', | 88 | 'General happiness with own health (0.0)', |
| 18 | 'Worry too long after embarrassment (2.0)', | 89 | 'Belief that own life is meaningful (0.0)', |
| 19 | 'Suffer from 'nerves' (0.0)', | 90 | 'Ever heard an un-real voice (0.0)', |
| 20 | 'Suffer from 'nerves' (2.0)', | 91 | 'Ever seen an un-real vision (0.0)', |
| 21 | 'Loneliness, isolation (0.0)', | 92 | 'Ever thought that life not worth living (0.0)', |
| 22 | 'Loneliness, isolation (2.0)', | 93 | 'Ever self-harmed (0.0)', |
| 23 | 'Guilty feelings (0.0)', | 94 | 'Ever contemplated self-harm (0.0)', |
| 24 | 'Guilty feelings (2.0)', | 95 | 'Felt hated by family member as a child (0.0)', |
| 25 | 'Risk taking (0.0)', | 96 | 'Physically abused by family as a child (0.0)', |
| 26 | 'Risk taking (2.0)', | 97 | 'Felt loved as a child (0.0)', |
| 27 | 'Frequency of depressed mood in last 2 weeks (0.0)', | 98 | 'Sexually molested as a child (0.0)', |
| 28 | 'Frequency of depressed mood in last 2 weeks (2.0)', | 99 | 'Someone to take to doctor when needed as a child (0.0)', |
| 29 | 'Frequency of unenthusiasm / disinterest in last 2 weeks (0.0)', | 100 | 'Felt irritable or had angry outbursts in past month (0.0)', |
| 30 | 'Frequency of unenthusiasm / disinterest in last 2 weeks (2.0)', | 101 | 'Avoided activities or situations because of previous stressful experience in past month (0.0)', |
| 31 | 'Frequency of tenseness / restlessness in last 2 weeks (0.0)', | 102 | 'Felt distant from other people in past month (0.0)', |
| 32 | 'Frequency of tenseness / restlessness in last 2 weeks (2.0)', | 103 | 'Repeated disturbing thoughts of stressful experience in past month (0.0)', |
| 33 | 'Frequency of tiredness / lethargy in last 2 weeks (0.0)', | 104 | 'Felt very upset when reminded of stressful experience in past month (0.0)', |
| 34 | 'Frequency of tiredness / lethargy in last 2 weeks (2.0)', | 105 | 'Ever sought or received professional help for mental distress (0.0)', |
| 35 | 'Seen doctor (GP) for nerves, anxiety, tension or depression (0.0)', | 106 | 'Ever suffered mental distress preventing usual activities (0.0)', |
| 36 | 'Seen doctor (GP) for nerves, anxiety, tension or depression (2.0)', | 107 | 'Ever had period of mania / excitability (0.0)', |
| 37 | 'Seen a psychiatrist for nerves, anxiety, tension or depression (0.0)', | 108 | 'Ever had period extreme irritability (0.0)', |
| 38 | 'Seen a psychiatrist for nerves, anxiety, tension or depression (2.0)', | 109 | 'Recent easy annoyance or irritability (0.0)', |
| 39 | 'Able to confide (0.0)', | 110 | 'Recent feelings or nervousness or anxiety (0.0)', |
| 40 | 'Able to confide (2.0)', | 111 | 'Recent feelings of inadequacy (0.0)', |
| 41 | 'Happiness (0.0)', | 112 | 'Recent trouble concentrating on things (0.0)', |
| 42 | 'Happiness (2.0)', | 113 | 'Recent inability to stop or control worrying (0.0)', |
| 43 | 'Work/job satisfaction (0.0)', | 114 | 'Recent feelings of depression (0.0)', |
| 44 | 'Work/job satisfaction (2.0)', | 115 | 'Recent poor appetite or overeating (0.0)', |
| 45 | 'Health satisfaction (0.0)', | 116 | 'Recent feelings of foreboding (0.0)', |
| 46 | 'Health satisfaction (2.0)', | 117 | 'Recent thoughts of suicide or self-harm (0.0)', |
| 47 | 'Family relationship satisfaction (0.0)', | 118 | 'Recent lack of interest or pleasure in doing things (0.0)', |
| 48 | 'Family relationship satisfaction (2.0)', | 119 | 'Recent trouble relaxing (0.0)', |
| 49 | 'Friendships satisfaction (0.0)', | 120 | 'Recent restlessness (0.0)', |
| 50 | 'Friendships satisfaction (2.0)', | 121 | 'Trouble falling or staying asleep, or sleeping too much (0.0)', |
| 51 | 'Financial situation satisfaction (0.0)', | 122 | 'Recent changes in speed/amount of moving or speaking (0.0)', |
| 52 | 'Financial situation satisfaction (2.0)', | 123 | 'Recent feelings of tiredness or low energy (0.0)', |
| 53 | 'Ever depressed for a whole week (0.0)', | 124 | 'Recent worrying too much about different things (0.0)', |
| 54 | 'Ever depressed for a whole week (2.0)', | 125 | 'Belittlement by partner or ex-partner as an adult (0.0)', |
| 55 | 'Longest period of depression (2.0)', | 126 | 'Been in a confiding relationship as an adult (0.0)', |
| 56 | 'Number of depression episodes (2.0)', | 127 | 'Physical violence by partner or ex-partner as an adult (0.0)', |
| 57 | 'Ever unenthusiastic/disinterested for a whole week (0.0)', | 128 | 'Sexual interference by partner or ex-partner without consent as an adult (0.0)', |
| 58 | 'Ever unenthusiastic/disinterested for a whole week (2.0)', | 129 | 'Able to pay rent/mortgage as an adult (0.0)', |
| 59 | 'Ever manic/hyper for 2 days (0.0)', | 130 | 'Been in serious accident believed to be life-threatening (0.0)', |
| 60 | 'Ever manic/hyper for 2 days (2.0)', | 131 | 'Been involved in combat or exposed to war-zone (0.0)', |
| 61 | 'Ever highly irritable/argumentative for 2 days (0.0)', | 132 | 'Diagnosed with life-threatening illness (0.0)', |
| 62 | 'Ever highly irritable/argumentative for 2 days (2.0)', | 133 | 'Victim of physically violent crime (0.0)', |
| 63 | 'Bipolar and major depression status (0.0)', | 134 | 'Witnessed sudden violent death (0.0)', |
| 64 | 'Neuroticism score (0.0)', | 135 | 'Victim of sexual assault (0.0)', |
| 65 | 'Ever addicted to any substance or behaviour (0.0)', | 136 | 'Did your sleep change? (0.0)', |
| 66 | 'Ever been injured or injured someone else through drinking alcohol (0.0)', | 137 | 'Trouble falling asleep (0.0)', |
| 67 | 'Frequency of inability to cease drinking in last year (0.0)', | 138 | 'Sleeping too much (0.0)', |
| 68 | 'Ever felt worried, tense, or anxious for most of a month or longer (0.0)', | 139 | 'Waking too early (0.0)', |
| 69 | 'Ever worried more than most people would in similar situation (0.0)', | 140 | 'Weight change during worst episode of depression (0.0)', |
| 70 | 'Age at first episode of depression (0.0)', | 141 | 'Ever been offered/sought treatment for anxiety (0.0)', |
| 71 | 'Age at last episode of depression (0.0)', | 142 | 'Ever been offered/sought treatment for depression (0.0)'] |

**Table S 4.**
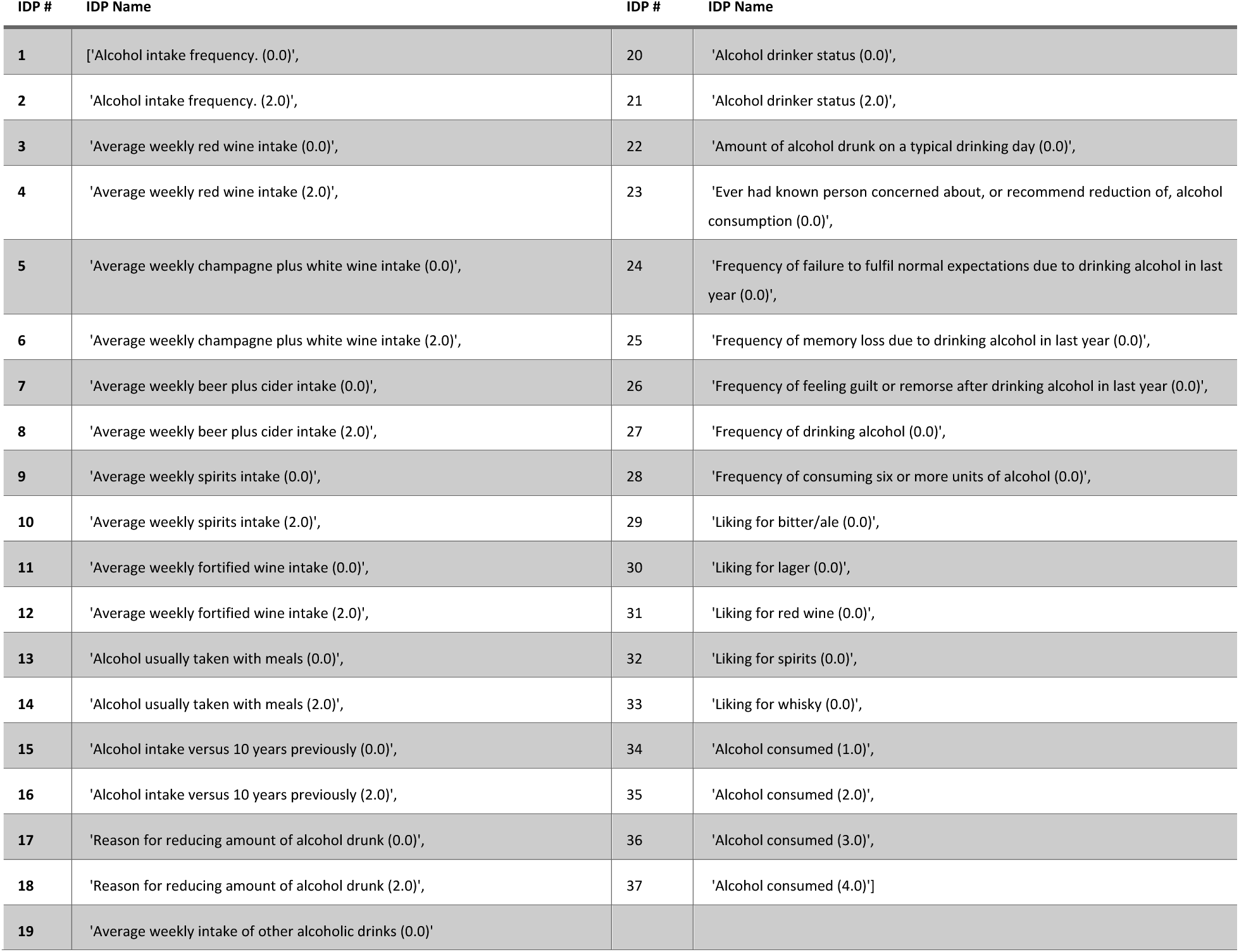
Alcohol nIDPs.

| IDP # | IDP Name | IDP # | IDP Name |
| --- | --- | --- | --- |
| 1 | ['Alcohol intake frequency. (0.0)', | 20 | 'Alcohol drinker status (0.0)', |
| 2 | 'Alcohol intake frequency. (2.0)', | 21 | 'Alcohol drinker status (2.0)', |
| 3 | 'Average weekly red wine intake (0.0)', | 22 | 'Amount of alcohol drunk on a typical drinking day (0.0)', |
| 4 | 'Average weekly red wine intake (2.0)', | 23 | 'Ever had known person concerned about, or recommend reduction of, alcohol consumption (0.0)', |
| 5 | 'Average weekly champagne plus white wine intake (0.0)', | 24 | 'Frequency of failure to fulfil normal expectations due to drinking alcohol in last year (0.0)', |
| 6 | 'Average weekly champagne plus white wine intake (2.0)', | 25 | 'Frequency of memory loss due to drinking alcohol in last year (0.0)', |
| 7 | 'Average weekly beer plus cider intake (0.0)', | 26 | 'Frequency of feeling guilt or remorse after drinking alcohol in last year (0.0)', |
| 8 | 'Average weekly beer plus cider intake (2.0)', | 27 | 'Frequency of drinking alcohol (0.0)', |
| 9 | 'Average weekly spirits intake (0.0)', | 28 | 'Frequency of consuming six or more units of alcohol (0.0)', |
| 10 | 'Average weekly spirits intake (2.0)', | 29 | 'Liking for bitter/ale (0.0)', |
| 11 | 'Average weekly fortified wine intake (0.0)', | 30 | 'Liking for lager (0.0)', |
| 12 | 'Average weekly fortified wine intake (2.0)', | 31 | 'Liking for red wine (0.0)', |
| 13 | 'Alcohol usually taken with meals (0.0)', | 32 | 'Liking for spirits (0.0)', |
| 14 | 'Alcohol usually taken with meals (2.0)', | 33 | 'Liking for whisky (0.0)', |
| 15 | 'Alcohol intake versus 10 years previously (0.0)', | 34 | 'Alcohol consumed (1.0)', |
| 16 | 'Alcohol intake versus 10 years previously (2.0)', | 35 | 'Alcohol consumed (2.0)', |
| 17 | 'Reason for reducing amount of alcohol drunk (0.0)', | 36 | 'Alcohol consumed (3.0)', |
| 18 | 'Reason for reducing amount of alcohol drunk (2.0)', | 37 | 'Alcohol consumed (4.0)'] |
| 19 | 'Average weekly intake of other alcoholic drinks (0.0)' |  |  |

**Table S 5.** Tobacco nIDPs.

| IDP # | IDP Name |
| --- | --- |
| 1 | 'Current tobacco smoking (0.0)', |
| 2 | 'Current tobacco smoking (2.0)', |
| 3 | 'Past tobacco smoking (0.0)', |
| 4 | 'Past tobacco smoking (2.0)', |
| 5 | 'Smoking/smokers in household (0.0)', |
| 6 | 'Smoking/smokers in household (2.0)', |
| 7 | 'Exposure to tobacco smoke at home (0.0)', |
| 8 | 'Exposure to tobacco smoke at home (2.0)', |
| 9 | 'Exposure to tobacco smoke outside home (0.0)', |
| 10 | 'Exposure to tobacco smoke outside home (2.0)', |
| 11 | 'Light smokers, at least 100 smokes in lifetime (0.0)', |
| 12 | 'Light smokers, at least 100 smokes in lifetime (2.0)', |
| 13 | 'Smoking status (0.0)', |
| 14 | 'Smoking status (2.0)', |
| 15 | 'Ever smoked (0.0)', |
| 16 | 'Ever smoked (2.0)', |
| 17 | 'Liking for cigarette smoking (0.0)', |
| 18 | 'Tobacco smoking (0.0)', |
| 19 | 'Amount of tobacco currently smoked (0.0)' |

**Table S 6.** Cardiovascular nIDPs.

| nIDP # | nIDP Name | nIDP # | nIDP Name |
| --- | --- | --- | --- |
| 1 | Pulse rate, automated reading (0.0), | 39 | Stroke volume during PWA (2.1), |
| 2 | Pulse rate, automated reading (0.1), | 40 | Mean arterial pressure during PWA (2.0), |
| 3 | Pulse rate, automated reading (2.0), | 41 | Mean arterial pressure during PWA (2.1), |
| 4 | Pulse rate, automated reading (2.1), | 42 | Ventricular rate (2.0), |
| 5 | Diastolic blood pressure, automated reading (0.0), | 43 | P duration (2.0), |
| 6 | Diastolic blood pressure, automated reading (0.1), | 44 | QRS duration (2.0), |
| 7 | Diastolic blood pressure, automated reading (2.0), | 45 | Systolic brachial blood pressure (2.0), |
| 8 | Diastolic blood pressure, automated reading (2.1), | 46 | Diastolic brachial blood pressure (2.0), |
| 9 | Systolic blood pressure, automated reading (0.0), | 47 | Cardiac index during PWA (2.0), |
| 10 | Systolic blood pressure, automated reading (0.1), | 48 | Cardiac index during PWA (2.1), |
| 11 | Systolic blood pressure, automated reading (2.0), | 49 | Pulse wave Arterial Stiffness index (0.0), |
| 12 | Systolic blood pressure, automated reading (2.1), | 50 | Pulse wave Arterial Stiffness index (2.0), |
| 13 | Pulse rate (0.0), | 51 | PQ interval (2.0), |
| 14 | Pulse rate (2.0), | 52 | QT interval (2.0), |
| 15 | Pulse wave reflection index (0.0), | 53 | QTC interval (2.0), |
| 16 | Pulse wave reflection index (2.0), | 54 | RR interval (2.0), |
| 17 | Pulse wave peak to peak time (0.0), | 55 | PP interval (2.0), |
| 18 | Pulse wave peak to peak time (2.0), | 56 | P axis (2.0), |
| 19 | Heart rate during PWA (2.0), | 57 | R axis (2.0), |
| 20 | Heart rate during PWA (2.1), | 58 | T axis (2.0), |
| 21 | Peripheral pulse pressure during PWA (2.0), | 59 | QRS num (2.0), |
| 22 | Central systolic blood pressure during PWA (2.0), | 60 | LV ejection fraction (2.0), |
| 23 | Central pulse pressure during PWA (2.0), | 61 | LV end diastolic volume (2.0), |
| 24 | Number of beats in waveform average for PWA (2.0), | 62 | LV end systolic volume (2.0), |
| 25 | Number of beats in waveform average for PWA (2.1), | 63 | LV stroke volume (2.0), |
| 26 | Central augmentation pressure during PWA (2.0), | 64 | Cardiac output (2.0), |
| 27 | Central augmentation pressure during PWA (2.1), | 65 | Average heart rate (2.0), |
| 28 | Augmentation index for PWA (2.0), | 66 | Minimum carotid IMT (intima-medial thickness) at 120 degrees (2.0), |
| 29 | Augmentation index for PWA (2.1), | 67 | Mean carotid IMT (intima-medial thickness) at 120 degrees (2.0), |
| 30 | Cardiac output during PWA (2.0), | 68 | Maximum carotid IMT (intima-medial thickness) at 120 degrees (2.0), |
| 31 | Cardiac output during PWA (2.1), | 69 | Minimum carotid IMT (intima-medial thickness) at 150 degrees (2.0), |
| 32 | End systolic pressure during PWA (2.0), | 70 | Mean carotid IMT (intima-medial thickness) at 150 degrees (2.0), |
| 33 | End systolic pressure during PWA (2.1), | 71 | Maximum carotid IMT (intima-medial thickness) at 150 degrees (2.0), |
| 34 | End systolic pressure index during PWA (2.0), | 72 | Minimum carotid IMT (intima-medial thickness) at 210 degrees (2.0), |
| 35 | End systolic pressure index during PWA (2.1), | 73 | Mean carotid IMT (intima-medial thickness) at 210 degrees (2.0), |
| 36 | Total peripheral resistance during PWA (2.0), | 74 | Maximum carotid IMT (intima-medial thickness) at 210 degrees (2.0), |
| 37 | Total peripheral resistance during PWA (2.1), | 75 | Minimum carotid IMT (intima-medial thickness) at 240 degrees (2.0), |
| 38 | Stroke volume during PWA (2.0), | 76 | Mean carotid IMT (intima-medial thickness) at 240 degrees (2.0), |
|  |  | 77 | Maximum carotid IMT (intima-medial thickness) at 240 degrees (2.0) |

**Table S 7.** Bone nIDPs.

| IDP # | IDP Name | IDP # | IDP Name |
| --- | --- | --- | --- |
| 1 | 'Heel bone mineral density (BMD) T-score, automated (0.0)', | 48 | 'Total BMD (bone mineral density) (2.0)', |
| 2 | 'Ankle spacing width (0.0)', | 49 | 'Total BMD (bone mineral density) T-score (2.0)', |
| 3 | 'Heel Broadband ultrasound attenuation, direct entry (0.0)', | 50 | 'Trunk BMC (bone mineral content) (2.0)', |
| 4 | 'Speed of sound through heel (0.0)', | 51 | 'Trunk BMD (bone mineral density) (2.0)', |
| 5 | 'Heel ultrasound method (right) (2.0)', | 52 | 'Femur shaft BMD (bone mineral density) (left) (2.0)', |
| 6 | 'Ankle spacing width (left) (0.0)', | 53 | 'Femur total BMD (bone mineral density) (left) (2.0)', |
| 7 | 'Ankle spacing width (left) (2.0)', | 54 | 'Femur total BMD (bone mineral density) T-score (left) (2.0)', |
| 8 | 'Heel broadband ultrasound attenuation (left) (0.0)', | 55 | 'Femur troch BMD (bone mineral density) (left) (2.0)', |
| 9 | 'Heel broadband ultrasound attenuation (left) (2.0)', | 56 | 'Femur wards BMD (bone mineral density) (left) (2.0)', |
| 10 | 'Speed of sound through heel (left) (0.0)', | 57 | 'Femur troch BMD (bone mineral density) T-score (left) (2.0)', |
| 11 | 'Speed of sound through heel (left) (2.0)', | 58 | 'Femur neck BMD (bone mineral density) (left) (2.0)', |
| 12 | 'Heel quantitative ultrasound index (QUI), direct entry (left) (0.0)', | 59 | 'Femur neck BMD (bone mineral density) T-score (left) (2.0)', |
| 13 | 'Heel quantitative ultrasound index (QUI), direct entry (left) (2.0)', | 60 | 'Femur wards BMD (bone mineral density) T-score (left) (2.0)', |
| 14 | 'Ankle spacing width (right) (0.0)', | 61 | 'Trunk bone area (2.0)', |
| 15 | 'Ankle spacing width (right) (2.0)', | 62 | 'Head bone area (2.0)', |
| 16 | 'Heel broadband ultrasound attenuation (right) (0.0)', | 63 | 'Head BMC (bone mineral content) (2.0)', |
| 17 | 'Heel broadband ultrasound attenuation (right) (2.0)', | 64 | 'Pelvis bone area (2.0)', |
| 18 | 'Speed of sound through heel (right) (0.0)', | 65 | 'Pelvis BMC (bone mineral content) (2.0)', |
| 19 | 'Speed of sound through heel (right) (2.0)', | 66 | 'Ribs bone area (2.0)', |
| 20 | 'Heel quantitative ultrasound index (QUI), direct entry (right) (0.0)', | 67 | 'Ribs BMC (bone mineral content) (2.0)', |
| 21 | 'Heel quantitative ultrasound index (QUI), direct entry (right) (2.0)', | 68 | 'Spine bone area (2.0)', |
| 22 | 'L1-L4 TBS (trabecular bone score) (2.0)', | 69 | 'Spine BMC (bone mineral content) (2.0)', |
| 23 | 'L1-L4 area (2.0)', | 70 | 'Arm bone area (left) (2.0)', |
| 24 | 'L1-L4 average height (2.0)', | 71 | 'Arm bone area (right) (2.0)', |
| 25 | 'L1-L4 average width (2.0)', | 72 | 'Arms combined bone area (2.0)', |
| 26 | 'L1-L4 BMC (bone mineral content) (2.0)', | 73 | 'Legs combined bone area (2.0)', |
| 27 | 'L1-L4 BMD (bone mineral density) (2.0)', | 74 | 'Femur neck bone area (left) (2.0)', |
| 28 | 'Femur neck BMD (bone mineral density) (right) (2.0)', | 75 | 'Femur neck bone area (right) (2.0)', |
| 29 | 'Femur neck BMD (bone mineral density) T-score (right) (2.0)', | 76 | 'Femur neck BMC (bone mineral content) (left) (2.0)', |
| 30 | 'Femur shaft BMD (bone mineral density) (right) (2.0)', | 77 | 'Femur neck BMC (bone mineral content) (right) (2.0)', |
| 31 | 'Femur total BMD (bone mineral density) (right) (2.0)', | 78 | 'Femur shaft bone area (left) (2.0)', |
| 32 | 'Femur total BMD (bone mineral density) T-score (right) (2.0)', | 79 | 'Femur shaft bone area (right) (2.0)', |
| 33 | 'Femur troch BMD (bone mineral density) (right) (2.0)', | 80 | 'Femur shaft BMC (bone mineral content) (left) (2.0)', |
| 34 | 'Femur troch BMD (bone mineral density) T-score (right) (2.0)', | 81 | 'Femur shaft BMC (bone mineral content) (right) (2.0)', |
| 35 | 'Femur wards BMD (bone mineral density) (right) (2.0)', | 82 | 'Femur total area (left) (2.0)', |
| 36 | 'Femur wards BMD (bone mineral density) T-score (right) (2.0)', | 83 | 'Femur total area (right) (2.0)', |
| 37 | 'Arm BMD (bone mineral density) (left) (2.0)', | 84 | 'Femur total BMC (bone mineral content) (left) (2.0)', |
| 38 | 'Arm BMD (bone mineral density) (right) (2.0)', | 85 | 'Femur total BMC (bone mineral content) (right) (2.0)', |
| 39 | 'Arms BMC (bone mineral content) (2.0)', | 86 | 'Femur troch bone area (left) (2.0)', |
| 40 | 'Arms BMD (bone mineral density) (2.0)', | 87 | 'Femur troch bone area (right) (2.0)', |
| 41 | 'Head BMD (bone mineral density) (2.0)', | 88 | 'Femur troch BMC (bone mineral content) (left) (2.0)', |
| 42 | 'Legs BMC (bone mineral content) (2.0)', | 89 | 'Femur troch BMC (bone mineral content) (right) (2.0)', |
| 43 | 'Legs BMD (bone mineral density) (2.0)', | 90 | 'Femur wards bone area (left) (2.0)', |
| 44 | 'Pelvis BMD (bone mineral density) (2.0)', | 91 | 'Femur wards bone area (right) (2.0)', |
| 45 | 'Ribs BMD (bone mineral density) (2.0)', | 92 | 'Femur wards BMC (bone mineral content) (left) (2.0)', |
| 46 | 'Spine BMD (bone mineral density) (2.0)', | 93 | 'Femur wards BMC (bone mineral content) (right) (2.0)'] |
| 47 | 'Total BMC (bone mineral content) (2.0)' |  |  |

